# eSCAN: Scan Regulatory Regions for Aggregate Association Testing using Whole Genome Sequencing Data

**DOI:** 10.1101/2020.11.30.405266

**Authors:** Yingxi Yang, Yuchen Yang, Le Huang, Jai G. Broome, Adolfo Correa, Alexander Reiner, NHLBI Trans-Omics for Precision Medicine (TOPMed) Consortium, Laura M. Raffield, Yun Li

**Author notes:** These authors contributed equally to this work (Yingxi Yang, Yuchen Yang, Laura M. Raffield, Yun Li). Corresponding author: Yun Li.

## Abstract

Multiple statistical methods for aggregate association testing have been developed for whole genome sequencing (WGS) data. Many aggregate variants in a given genomic window and ignore existing knowledge to define test regions, resulting in many identified regions not clearly linked to genes and thus limiting biological understanding. Functional information from new technologies (such as Hi-C and its derivatives), which can help link enhancers to their effector genes, can be leveraged to predefine variant sets for aggregate testing in WGS data. Here we propose the eSCAN (Scan the Enhancers) method for genome-wide assessment of enhancer regions in sequencing studies, combining the advantages of dynamic window selection in SCANG, a previously developed method, with the advantages of incorporating putative regulatory regions from annotation. eSCAN, by searching in putative enhancer, increases statistical power and aids mechanistic interpretation, as demonstrated by extensive simulation studies. We also apply eSCAN for blood cell traits using TOPMed WGS data. Results from real data analysis show that eSCAN is able to capture more significant signals, and these signals are of shorter length (indicating higher resolution fine-mapping capability) and drive association of larger regions detected by other methods.

## Introduction

In genome-wide association studies (GWAS), most significantly associated variants are located outside coding regions of genes, making it difficult to interpret the biological function of associated variants. Statistical power to detect rare variant associations in noncoding regions, which is of increasing importance with the advent of large-scale whole genome sequencing (WGS) studies, is also limited with a standard single variant GWAS approach. Aggregate testing is necessary to increase statistical power to detect rare variant associations; linking noncoding variants to their likely effector genes is necessary for interpretation of identified aggregate signals. Many standard methods for aggregate analysis of the noncoding genome are agnostic to regulatory and functional annotation (for example, standard sliding window analysis, where all variants in a given location bin (for example a 5 kb or 10 kb window) are analyzed, followed by analysis of a subsequent partially overlapping window, until each chromosome is assessed in full)(1-3). SCANG has recently been proposed as an improvement on conventional sliding-window procedures, with the ability to detect the existence and locations of association regions with increased statistical power (4). SCANG allows sliding-windows to have different sizes within a pre-specified range and then searches all the possible windows across the genome, increasing statistical power. However, since SCANG tests all possible windows, it can “randomly” identify some regions across the genome regardless of their biological functions. Identified regions could often cross multiple enhancer regions with distinct functions, thus impeding the identification of biologically important enhancers and their target genes. This cross-boundary issue may also lead to a higher false positive rate in a fine-mapping sense. The whole region/chromosome in which the detected regions are located may not be a false positive, but locations of the detected regions will not match the true association regions. Moreover, SCANG applies SKAT to all candidate windows, but computing p-values in SKAT requires eigen decomposition (5). This analysis method is therefore very time-consuming and has high computational costs, which may not be feasible for increasingly large genome-wide studies.

In addition to sliding window approaches, many analyses of WGS data rely on aggregate tests of predefined variant sets, attempting to link the most likely regulatory variants (as defined by tissue specific histone marks, open chromatin data, sequence conservation, etc) to genes prior to association testing, with variants assigned to genes based on either physical proximity or chromatin conformation (1,3). There is increasing data available to define these tissue specific regulatory regions, which are known to show enrichment for GWAS identified noncoding variant signals (6-8). Recent biotechnological advances based on Chromatin Conformation Capture (3C), such as promoter capture Hi-C data, can also better link gene promoters to enhancers based on their physical interactions in 3D space (9). We here propose an extension of SCANG which combines the advantages of both scanning and fixed variant set methods (see **Fig. 1** for illustration). Our eSCAN (or “scan the enhancers” with “enhancers” as a shorthand for any potential regulatory regions in the genome) method can integrate various types of functional information, including chromatin accessibility, histone markers, and 3D chromatin conformation. There can be a significant distance between a gene and its regulatory regions; simply expanding the size of the window to include kilobases of genomic data around each gene will include too many non-causal SNPs, giving rise to power loss, as well as difficulties in results interpretation (9). Our proposed framework can enhance statistical power for identifying new regions of association in the noncoding genome. We particularly focus on integration of 3D spatial information, which has not yet been fully exploited in most WGS association testing studies. Our method allows users to input broadly defined regulatory/enhancer regions and then select those which are most likely relevant to a given phenotype, in a statistically powerful framework.

**Figure 1:**
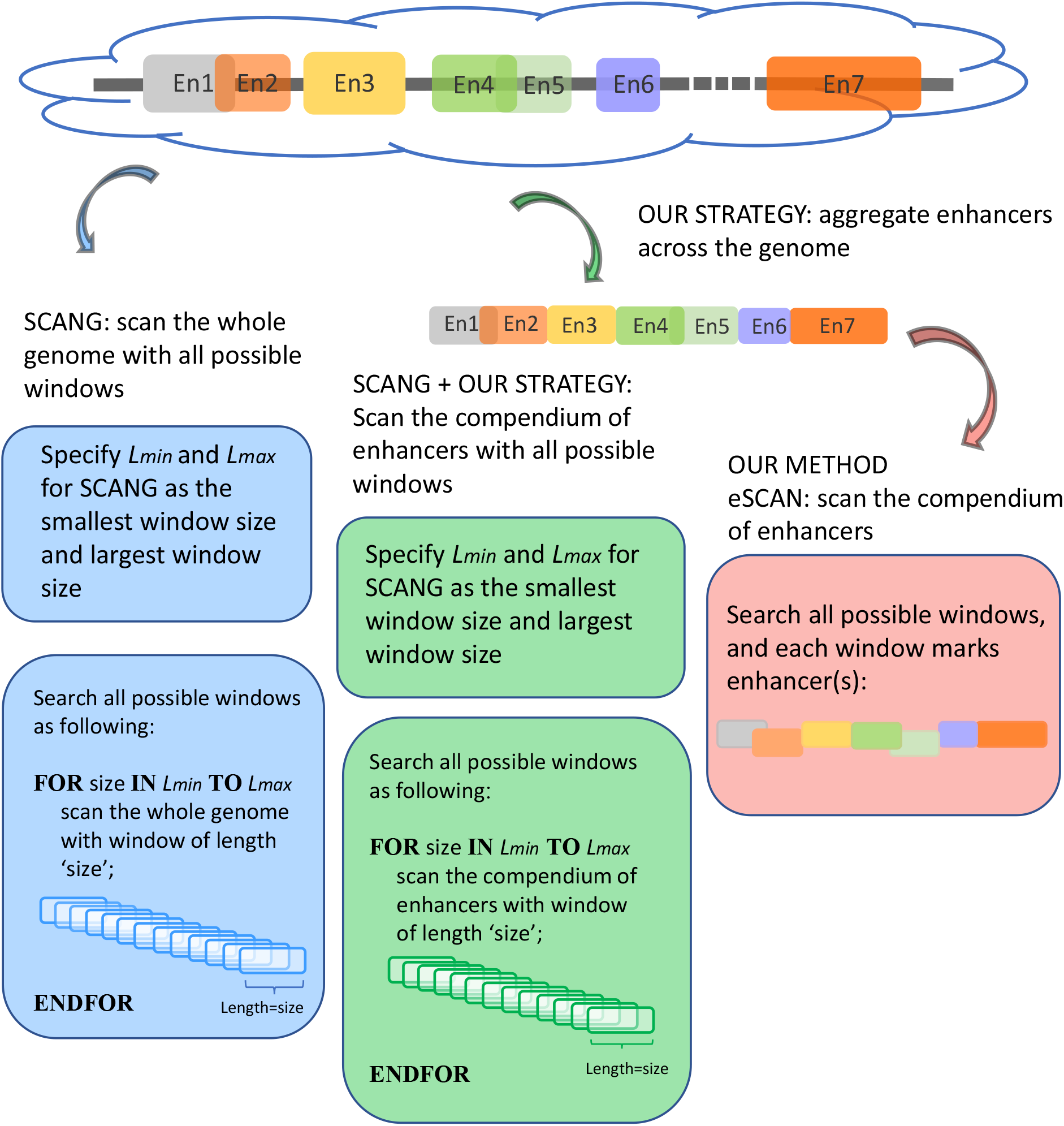
An illustration of eSCAN.

Given our incomplete understanding of chromatin conformation and enhancer annotation, an annotation agnostic approach such as SCANG does have some advantages, in that no prior information is needed for rare variant testing. However, our simulations and the real data example presented here demonstrate the advantages of our eSCAN method, which can flexibly accommodate multiple types of annotation information and shows significant power gains over SCANG, as well as a lower false positive rate in different scenarios for both continuous and dichotomous traits. These advantages are demonstrated in our application of eSCAN to TOPMed WGS analyses of four blood cell traits in the Women’s Health Initiative (WHI) study, with replication in Jackson Heart Study (JHS).

## Materials and Methods

### eSCAN Framework

#### Overview of eSCAN

The eSCAN procedure can be split into two steps. First is the enhancer-screening step, where set-based *p*-values for each enhancer are calculated by fastSKAT utilizing different weights, then p-values are combined by the Cauchy method via aggregated Cauchy association test (ACAT) (4). eSCAN then defines potential significant enhancers using estimated significance threshold, either an empirical estimation based on Monte Carlo simulation or an analytical estimation by extreme value distribution (10). Second, eSCAN performs a dynamic sliding-window scanning within each of the potential significant enhancers to further narrow down the associated region.

#### Step A: One Set-based *p*-value for Each Enhancer by Omnibus FastSKAT

eSCAN considers each putative enhancer region as a searching window and first calculates *p*-values for each window using fastSKAT. FastSKAT applies randomized singular value decomposition (SVD) to rapidly analyze much larger regions than standard SKAT (11), which makes it computationally feasible to deal with long super enhancer regions.

FastSKAT calculates the test statistics *Q* in the same way SKAT does (11). It differs from the standard SKAT test in its approximation of the null distribution of *Q*, using the basic Satterthwaite approximation with an additional remainder term,

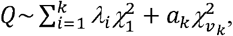

where *λ*_1_, *λ*_2_,…, *λ*_k_ are the largest *k* eigenvalues of the covariance matrix of the genotypes. And the scaling and degree of freedom in the remainder term is obtained by moment-matching

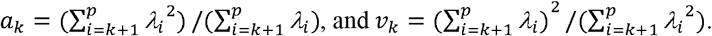

FastSKAT requires only k leading eigenvalues rather than full eigenvalues. And the leading eigenvalues calculation is implemented by the random projection approach, which powerfully combines probability and matrix theory. The computation can be split into the following two stages. The first stage is dimension reduction, constructing a new matrix whose rank is lower than the input matrix but accurately approximates its range. The second stage is to implement a standard factorization, such as QR and SVD, of the dimension reduced matrix obtained from the first stage.

For the choice of weights when calculating the test statistic *Q*, following SCANG, we adopted two commonly used weights, Beta (1,1) and Beta (1,25). Beta (1,1) corresponds to equal weights for all variants while Beta (1,25) results in up-weighting rarer variants based on the assumption that rarer variants tend to exert larger effects (12). A set-based *p* value is obtained by combining *p*-values from the above two sets of weights using the Cauchy method via ACAT (13):

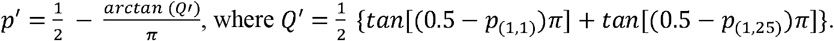

where *p*_(1,1)_ and *p*_(1,2, 5)_ denote the *p*-values of fastSKAT using *a*_1_ *a*_2_=1 and *a*_1_, *a*_2_ =25 in the beta distribution density function.

#### Step B: Empirically or Analytically Estimating the Significance Threshold

After obtaining a single *p*-value for each enhancer region, eSCAN next computes the significance threshold. Frequent physical overlaps between enhancers as well as linkage disequilibrium (LD) among variants across enhancers in sequencing data tend to elicit high correlation between the *p*-values of these enhancers, making the classic Bonferroni correction for multiple testing too conservative, leading to power loss. Therefore, we consider alternative methods to estimate the significance threshold. Specifically, we provide both an empirical and an analytical approach to derive the significance threshold.

The classic Monte Carlo method based on a common distribution of test statistics is inappropriate given that the mixture of chi-square distributions in fastSKAT is a set-based one, which means the test statistics in different enhancers would follow different null distributions. Hence, we adopt a Monte Carlo method on the basis of the common distribution of *p* values, which is similar to that in the SCANG paper (4). We present the following statistical framework to estimate the threshold.

1. An *n* x 1 pseudo-residual vector 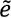 is generated from a multivariate normal distribution *N* (0, I_n_).
2. A *p* x 1 pseudo-score vector *Ũ* is calculated by 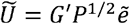, where *G* is the *n* x *p* genotype matrix; n is sample size; p is the number of rare variants across all enhancers in sequencing data. 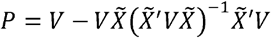 is the projection matrix defined in the SKAT paper(5), where 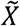 is a design matrix containing the intercept term. When the phenotype is continuous, 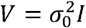, where 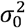 is the estimate of the variance of error term under the global null hypothesis. When the phenotype is dichotomous 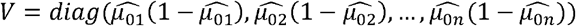 where 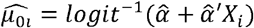. Note that although pseudo-scores differ across iterations, they follow the same distribution *N*(0, *G*’*PG*).
3. Given the pseudo-score *u* (*Ũ*_1_, *Ũ*_2_,…, Ũ_*p*_,) we calculate single set-based *p* value p for each enhancer by running the omnibus test, as described in step A above. For the computational formula of *Q*, simply substitute *Ũ*_*j*_ for U_*j*_. It is worth noting that although the individual score statistic *Q* alculated from an observed phenotype might not be normally distributed, the set-based test statistics *Q* follows the same distribution as 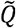 obtained from pseudo-score does. Consequently, pseudo *p* value *p* and observed 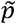 value share the same distribution as well.
4. Take the minimum 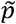_*min*_ among the pseudo *p* values 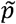 for each enhancer.
5. Repeat step 1-4 B times; get 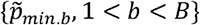. Then, to control the genome-wide type I error rate at the α level, we choose as the empirical threshold the *α*^*th*^ quantile of *B* 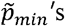 where B is the number of iterations.

We can then select the enhancers whose observed *p*-values are below the empirical threshold and define them as detected potentially causal enhancers.

In addition, we provide an analytical estimation of the significance threshold using the Gumbel distribution, which is also based on the common distribution of *p*-values following the WGScan method (10). In order to estimate the parameters of the Gumbel distribution, a resampling approach is still needed. The difference is that here the resampling is used to estimate the first and second moments of the Gumbel distribution. In practice, we implement the following step instead of the step 5 above.

5*. Repeat step 1-4 *B* times; get 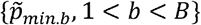. Use their sample mean and variance to approximate the first and second moments of the Gumbel distribution.

Then calculate 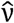 and 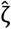 by *E*(*x*)*ν ζ γ* and 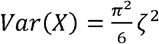, where *γ* ≈ 0.57721 isbthe Euler-Mascheroni constant. Calculate significance threshold 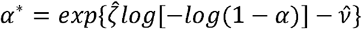.

### Simulations under the Null Model

We first evaluate the performance of eSCAN under the null model. The sequencing data used in our simulations is provided in the SCANG package, where 20,000 chromosomes for a 5 Mb region (representing the whole genome, in the interest of computational efficiency) were simulated using COSI, leveraging its calibrated model to closely resemble the LD patterns from African Americans (4). Only rare variants whose MAF is below 0.05 were used for both eSCAN and SCANG analysis. For each simulation, 400 enhancers are randomly generated with length of 3, 4 or 5 kb (each has a probability of 1/3) across the genome where the enhancers are allowed to overlap with each other. On average, each simulated enhancer had a length of 4025 bp and contained 122 variants with MAF below the pre-specified threshold, 5%.

We simulated continuous/dichotomous phenotype data using the following models:

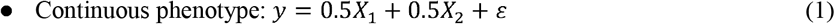

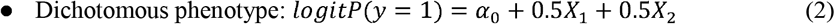

where *X*_1_ is a continuous covariate simulated from a standard normal distribution, *X*_2_ is a binary covariate generated from a Bernoulli distribution with *p* = 0.5, and c is an error term following a standard normal distribution, and *α*_0_ is a parameter to set the prevalence to 1%.

For both continuous and *α*_0_ dichotomous simulations, we applied eSCAN to 1,000 replicates with sample sizes of 2,500, 5,000 and 10,000, respectively, and set the genome-wide type I error rate at 0.05. The empirical type I error rate was estimated by the proportion of rejections under the null where a rejection was declared if eSCAN reported at least one enhancer as significant.

### Simulations under the Alternative Model

To assess eSCAN under the alternative model, i.e. power and false positive rate, 10% of enhancers were randomly selected as causal from the 400 simulated enhancer regions. Within each causal enhancer 20% of variants were randomly chosen as causal variants with effect sizes β’s whose distributional specification is provided below (**Fig. S1**). Then we used these causal variants to create phenotypes together with the covariates described above:

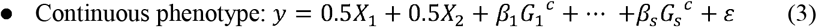

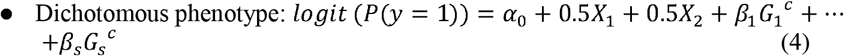

where *G*_1_^*c*^, *G*_2_^*c*^, …, *G*_*S*_^*c*^are the genotypes of the *s* causal rare variants in the causal enhancers. *β*’s are effect sizes of the causal variants. *α*_0_, *X*_1_, *X*_2_, and *c* remain the same as defined in equation (1) and (2). Based on the assumption that rarer variants tend to exert larger effects, for both traits we set *β*_*i*_ |*iog* _10_ *MAF*_*i*_ |, a decreasing function of MAF of the *i*-th variant, where is a parameter to control the magnitude of effect size. For continuous traits, 0.18, giving a *β* = 0.90 for variants with MAF =1 x 10^−5^and a smaller effect size *β* = 0.36 for less rare variants with MAF = 1 x 10^−5^; for dichotomous traits, *c =* 0.255, giving an OR = 3.579 for variants with MAF 1 x 10^−5^and a smaller OR = 1.665 for less rare variants with MAF = 1 x 10^−2^.

We then applied eSCAN and SCANG to each simulated scenario to benchmark theirVperformances in terms of power and false positive rate. In order to evaluate the value of aggregating variants residing in enhancer regions, we additionally applied an enhancer-based SCANG that scans the subset of variants which reside in enhancer regions. In both applications of SCANG, the ranges of sliding window size *L*_*min*_ and _*max*_ were set to be 10^*th*^ quantile and 90^*th*^ quantile of the empirical distribution of the number of rare variants within the enhancers, respectively. For other parameters such as skip-length of searching windows in the SCANG, we adopted the package default. We then compared eSCAN with the two SCANGs (default SCANG and enhancer-based SCANG), using four metrics (more details below), namely causal-variant detection rate, causal-enhancer detection rate, variant false positive rate and enhancer false positive rate (defined below).

We further evaluated eSCAN in more simulation scenarios to verify eSCAN’s robustness to the proportion of causal enhancers and the proportion of causal variants within the causal enhancers. We set the percentage of causal enhancers across the genome to be 5%, 10% and 15% respectively, with a fixed proportion of 20% causal variants within causal enhancers. We next conducted simulations for the percentage of causal variants being 10%, 15% and 20% respectively, where the proportion of causal enhancers was fixed to be 10%.

To evaluate the methods under alternative models, we here define four criteria to evaluate the performance of the methods under the alternative model. For power assessment, we adopted two criteria. As a power metric at variant level, *causal-variant detection rate* is calculated as the number of detected causal variants divided by the total number of causal variants, where a causal variant is deemed as detected if it is located in one of the detected enhancers or regions, similar to the power calculations performed in Li *et al*. (4). As an estimation of power at enhancer level, *causal-enhancer detection rate* is similarly calculated as the number of detected causal enhancers divided by the total number of causal enhancers. In eSCAN, the detected causal enhancers are the output directly obtained from our method, but this is not the case in SCANG, since the detected regions given by SCANG wouldn’t be expected to match the enhancer boundaries in an exact way. We here define a causal enhancer as detected if the overlapping fraction of the causal enhancer and one of the detected regions is larger than 0.5, where the overlapping fraction is the number of variants included in both the causal enhancer and the detected region, divided by the number of variants located in the causal enhancer.

Likewise, we used two criteria for the assessment of false positive rate, one at the variant level and the other at the enhancer level. The *variant false positive rate* is the number of false-identified variants divided by the total number of non-causal variants, where a variant is deemed as false-identified if it is located in one of the detected regions while in fact it does not reside within any causal enhancer. Similarly, the *enhancer false positive rate* is calculated as the number of false-identified enhancers over the total number of non-causal enhancers, where an enhancer is false-identified if it is detected but is not causal.

### Application to Blood Cell Traits using TOPMed WGS in JHS and WHI Studies

To assess the performance of eSCAN in real data, we applied eSCAN and enhancer-based SCANG for association analysis of white blood cell count (WBC), platelet count (PLT), hemoglobin (HGB), and hematocrit (HCT) using the WGS data in 1,970 unrelated individuals (defined as kinship coefficient < 0.2 with all other included individuals) from the Jackson Heart Study (JHS) and 10,727 unrelated individuals from the Women’s Health Initiative (WHI) studies. We consider only uncommon variants with MAF < 0.05. Jointly called genotypes from >30x WGS data were available for both cohorts through the TOPMed consortium; detailed methods are available at https://www.nhlbiwgs.org/topmed-whole-genome-sequencing-methods-freeze-8. The design of JHS (14) and WHI (15) has been described in detail previously. Exclusion criteria for phenotypic were WBC > 200 x 10^9^/L, HGB > 20g/dL, HCT > 60% and PLT > 1000 x 10^9^/L. We adjusted for the first 11 principal components (PCs) created by *pcair* function from the GENESIS R package, along with sex, age and square of age for JHS. For WHI, we used the same covariates, excluding sex (as the study includes only women). We additionally adjusted for the case/control status and, for WBC, the Duffy null variant rs2814778, a non-coding variant residing in gene *ACKR1*, Atypical Chemokine Receptor 1 in the Duffy blood group. This variant, common in individuals of African ancestry, explains 7-20% of the variation in WBC among African Americans (16,17). Because of its huge effect size, this variant is conventionally included as a covariate when performing association analysis with WBC among African Americans to avoid severe inflation of test statistics due to linkage disequilibrium (18).

For eSCAN, enhancers were defined using promoter capture-Hi-C (PC-HiC) data in white blood cell types (19). Specifically, we considered a genomic region as an enhancer if 1) it interacts with a promoter region identified by PC-HiC (bait region), or 2) it, although initially considered as a promoter candidate by PC-HiC (i.e., with some bait designed to cover the region), interacts with another gene’s promoter region, where the promoter region is defined as +/- 500 bp from the transcriptional starting site (TSS). Definition of enhancers with eSCAN is flexible and can be guided by regulatory region annotation and chromatin conformation data available for relevant cell types for a given trait. For enhancer-based SCANG, we implemented association tests for the subset of rare variants falling into any enhancer region as defined using PC-HiC annotation. For the default SCANG, due to the limited computational feasibility, we only performed the analysis for WBC. The range of searching window sizes was set by specifying the minimum and maximum numbers of variants in searching windows between *L*_*min*_ = 100, *L*_*max*_ = 180 for JHS and *L*_min_ = 140,*L*_*max*_ = 220 for WHI. For other parameters such as skip-length of searching windows in the SCANG, we adopted the package defaults.

## Results

### Simulation Results

We first evaluated the performance of eSCAN using simulated data under the null model. On average, each simulated enhancer had a length of 4025 bp and contained 122 variants with MAF below 5%. For both continuous and dichotomous simulations, we applied eSCAN to 1,000 replicates with sample sizes of 2,500, 5,000 and 10,000, respectively, and set the genome-wide type I error rate at 0.05. Under all scenarios, our method has a well-controlled genome-wide type I error rate (**Table 1**).

**Table 1:** Genome-wide empirical type I error rates of eSCAN from simulation studies, shown for different sample sizes and trait distributions.

| Sample size | n = 2,500 | n = 5,000 | n = 10,000 |
| --- | --- | --- | --- |
| Continuous Traits | 0.003 | 0.002 | 0.001 |
| Dichotomous Traits | 0.001 | 0.001 | 0.002 |

To assess eSCAN under the alternative model, we applied eSCAN and two SCANGs, i.e. the default SCANG and enhancer based SCANG, to a wide range of simulated scenario to benchmark their performances in terms of power and false positive rate, using four metrics described above. For continuous traits, both the enhancer-based SCANG and our eSCAN analysis showed higher power than the default SCANG, at both the variant level and the enhancer level (**Fig. 2a-b**), for all tested sample sizes, suggesting the benefit of aggregating variants using enhancer information. Notably, the power gain between eSCAN and enhancer-based SCANG is much more pronounced than that between enhancer-based SCANG and default SCANG. eSCAN increases the variant-level power by 23.50%, 45.94% and 27.98% for the three tested sample sizes, respectively; and boosts the enhancer-level power by 17.60%, 45.47% and 24.14%, respectively.

**Figure 2:**
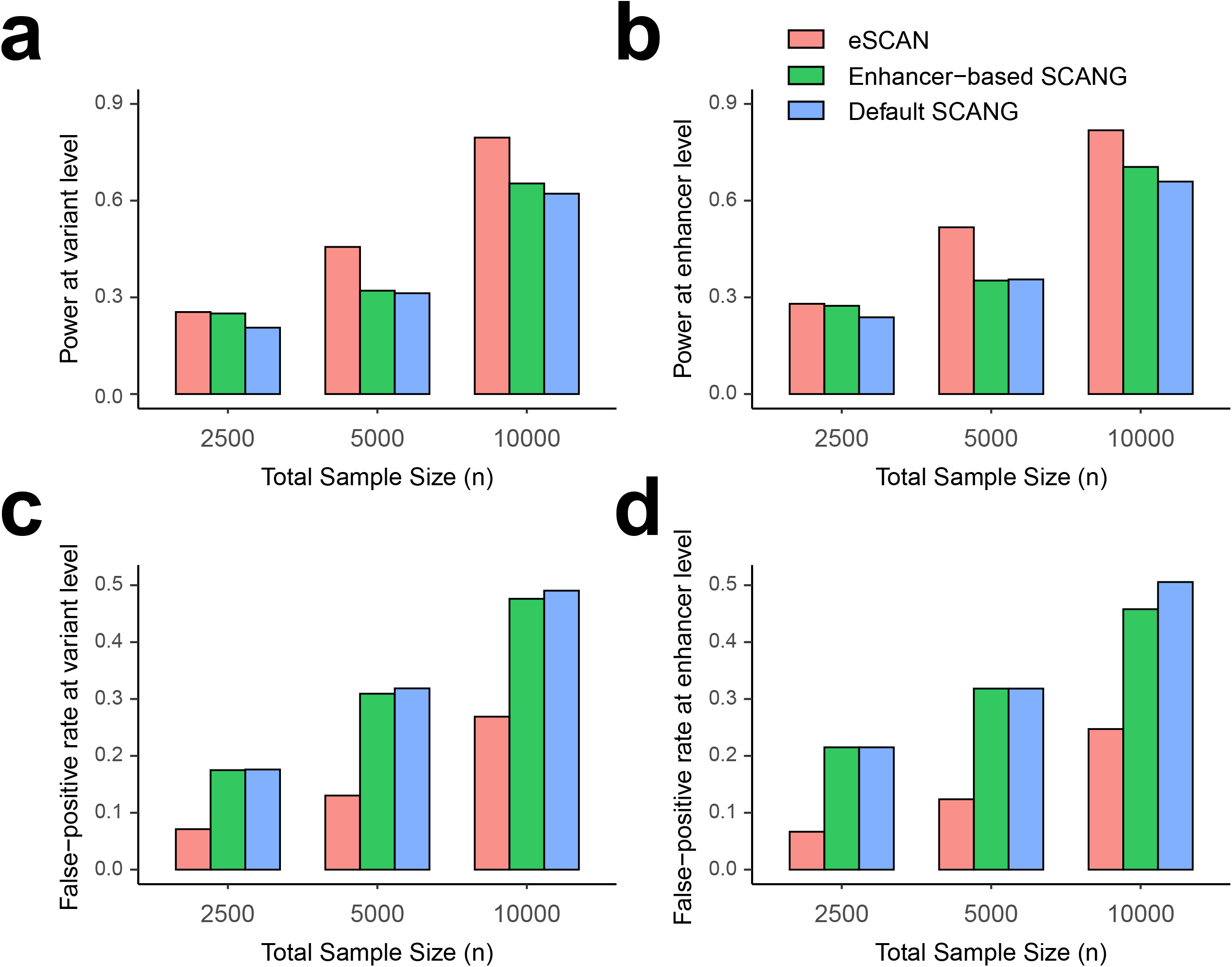
Power and false positive rate comparison of eSCAN and SCANG for continuous outcome at various sample sizes. We evaluated the performance of eSCAN for continuous outcome at various sample sizes. The total sample sizes considered were 2,500, 5,000 and 10,000. At each sample size, we compared three methods: eSCAN and two versions of SCANGs: enhancer-based SCANG (aggregating enhancers across the genome) and default SCANG (scan the whole genome). We evaluated power and false-positive rate at both variant level and enhancer level. **a**. Power at the variant level (a.k.a. causal variant detection rate). **b**. Power at enhancer level (a.k.a. causal enhancer detection rate). **c**. False positive rate at variant level. **d**. False positive rate at enhancer level.

With respect to false positive rate, eSCAN showed a remarkably lower false positive rate than those from the two SCANG procedures, at both the variant-level and enhancer-level (**Fig. 2c-d**). For the two SCANG procedures, the false positive rates are high because of the aforementioned cross-boundary issue accompanied with the scanning procedure. Although the power of SCANG increases as the sample size increases, indicating its ability to detect more causal enhancers when more individuals’ data is available, the false positive rate also increases dramatically (**Fig. 2c-d**). For a sample size of 10,000, the enhancer-level false positive rate of the default SCANG is up to 0.51. Numerically, as the sample size increases from 5,000 to 10,000, the increase rate of enhancer-level false positive rate is 47.99%, close to the power gain of 49.33%, suggesting results may become less trustworthy. By contrast, eSCAN reduces the false positive rate by 69%, 61% and 51% at the enhancer-level for all tested sample sizes, respectively. Similar reductions at the variant-level are observed by 60%, 59% and 45%. These results demonstrate eSCAN’s capabilities to powerfully and accurately detect causal enhancers. The results from dichotomous traits also show that eSCAN outperformed the two SCANG approaches (**Fig. S2**). We further evaluated eSCAN in more simulation scenarios to verify eSCAN’s robustness to the proportion of causal enhancers and the proportion of causal variants within the causal enhancers. Results show that these gains are robust to choice of parameters (**Fig. S2-4**), suggesting that the superiority of eSCAN is inherent and is not accidentally driven by the choices of parameters in the simulations.

### Real Data Results for Blood Cell Traits using TOPMed

To assess the performance of eSCAN in real data, we compared eSCAN to both enhancer-based SCANG and the default SCANG using WGS data in 10,727 discovery samples from the Women’s Health Initiative (WHI) and 1,970 replication samples from the Jackson Heart Study (JHS) (**Table S1**). We only considered variants with a minor allele frequency < 5% in each cohort. Windows with a total minor allele count (MAC) < 10 were excluded from the analysis. To achieve a fair comparison, we first applied eSCAN and enhancer-based SCANG for association analysis between putative enhancers and four blood cell traits measured at baseline in WHI, white blood cell count (WBC), hemoglobin (HGB), hematocrit (HCT) and platelets (PLT), with a genome-wide error rate at the level of 0.05 by Bonferroni correction in both methods. For eSCAN, enhancers were defined using promoter capture-Hi-C (PC-HiC) data in any tested white blood cell type (including neutrophils, monocytes, and lymphocytes) for WBC, erythroblasts for HGB and HCT and megakaryocytes for PLT(19), defining any noncoding region with statistically significant interactions with a gene promoter as an enhancer region. For enhancer-based SCANG, we analysed the subset of rare variants falling into any enhancer region as defined using PC-HiC annotation. For the default SCANG, due to the limited computational feasibility, we only performed the analysis for WBC.

Overall, eSCAN detected 19 significant regions associated with blood cell traits while enhancer-based SCANG only detected 7 regions (**Table 2**; **Table S2 and S3**; **Fig. S5** and **S6**). Also, eSCAN showed consistently smaller p-values for top regions compared with enhancer-based SCANG (**Fig. S5A-D; Table 2**). Among the 19 genome-wide significant regions detected by eSCAN in the unconditional analysis, 4 were located within +/- 500kb of known GWAS loci and were still significant at the Bonferroni correction level of 0.05/4 after conditioning on known blood cell trait GWAS loci (12,18,20-26) (**Table 2; Table S4**). Also, of the significant regions, two were replicated at 0.05 level in replication samples. Note that the low replication rate is likely due in large part to the much smaller sample size of the JHS replication cohort; we also note as a limitation that we did not correct for multiple testing in these replication analyses, due to this small sample size.

**Table 2:**
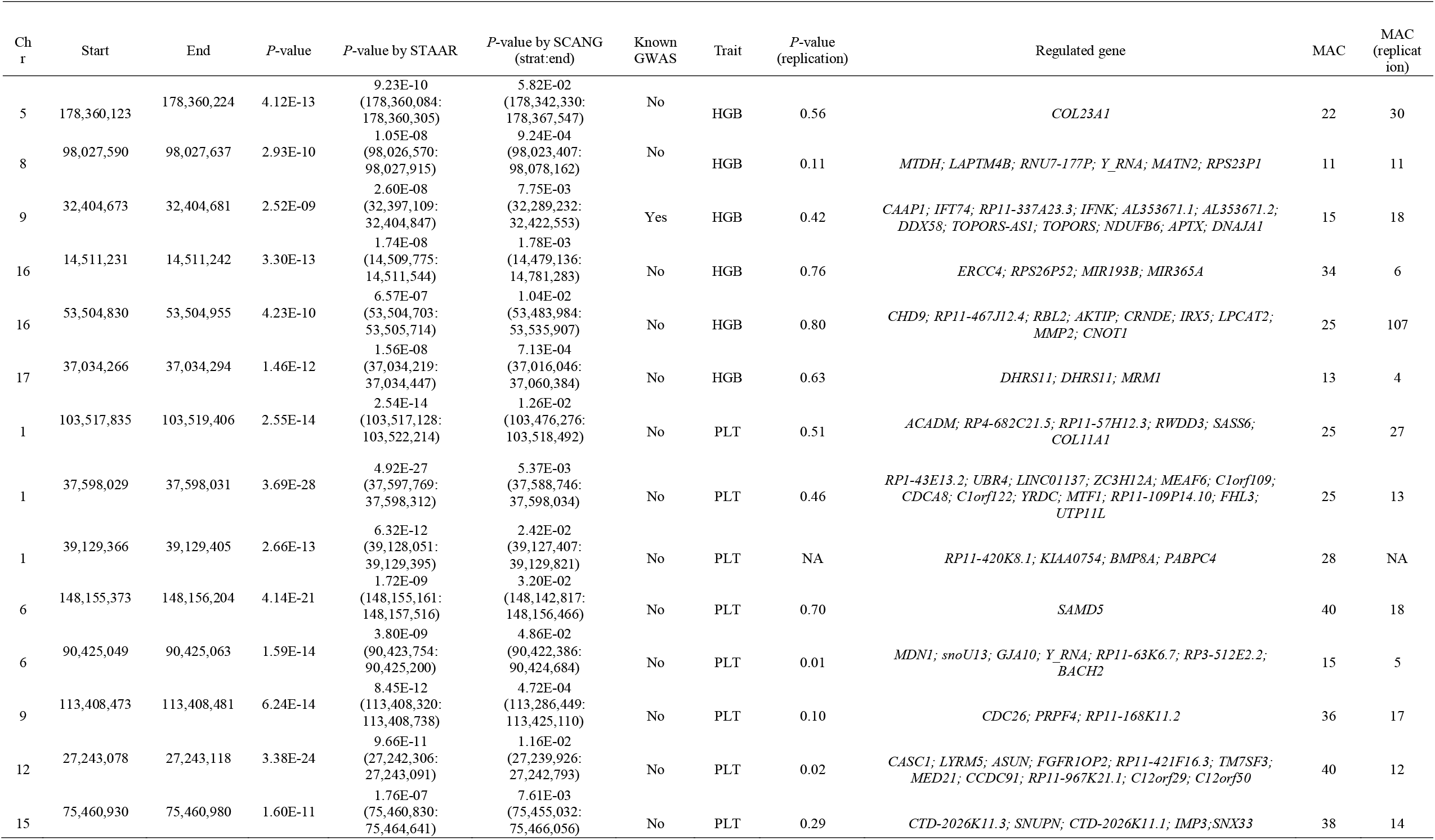
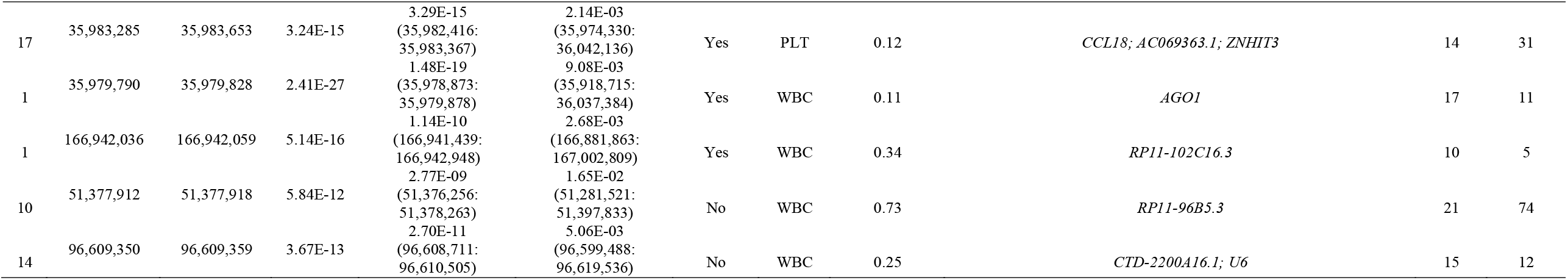
Significant results by eSCAN for blood cell traits in TOPMed whole genome sequencing data. Chromosome, start position (hg38), end position (hg38), *p*-value in discovery samples (Women’s Health Initiative, WHI), *p*-value of nearest region tested by enhancer-based SCANG, known GWAS loci within +/- 500kb, associated trait (HGB or hemoglobin, PLT or platelet count, WBC or white blood cell count), *p*-value in the replication samples (Jackson Heart Study, JHS), gene(s) regulated by the detected region, total minor allele count (MAC) in the discovery samples, total MAC in the replication samples.

To more comprehensively compare the top regions of eSCAN and two SCANG procedures, enhancer-based SCANG and default SCANG, we relaxed the significance level for WBC by using the empirical threshold. The detected regions by eSCAN are of shorter length and contains fewer variants than those identified by the two SCANG variants (**Fig. S7b**). Also, each region identified by eSCAN contains a single regulatory element based on annotation from promoter capture Hi-C. By contrast, regions identified by SCANG can cross multiple regulatory regions (**Fig. S7c**), which indicates that, with the help of enhancer information, eSCAN can more effectively narrow down variants and/or regulatory regions associated with a trait of interest than SCANG. We further investigated a segment on chr10 where two signals were detected by enhancer-based SCANG and four by eSCAN. The two regions from SCANG overlapped the four eSCAN signals. All four were smaller in size than the SCANG detected regions. We also note that each SCANG signal contains two eSCAN signals (**Fig. 3a-c**). We then removed the associated variants in the overlapped regions between eSCAN and SCANG (which are regions detected by eSCAN since in both cases, the eSCAN regions are subsets of the SCANG regions), and re-did SCANG analysis using the retained variants only. Both regions then became insignificant (*p*-values > 0.02) using SCANG (**Fig. 3d**), suggesting that the sub-regions detected by eSCAN were most likely the functional regions contributing to the original significant signal.

**Figure 3:**
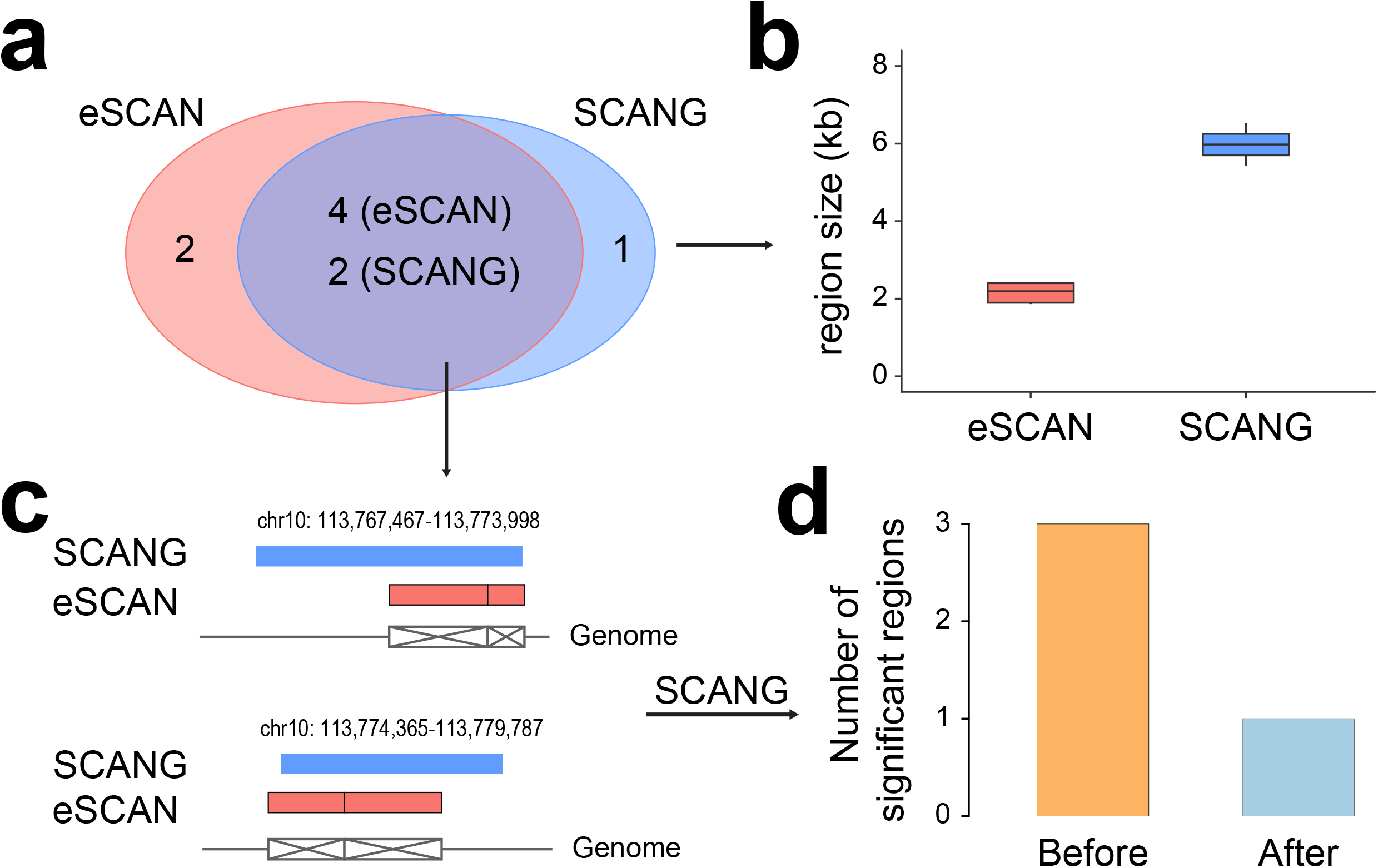
A segment on chr10 where two signals were detected by SCANG and four by eSCAN. We further investigated a segment on chr10 where two signals were detected by SCANG and four by eSCAN. The two regions from SCANG (chr10:113,767,467-113,773,998 with *p*-value = 2.66×10^−7^ and chr10:113,774,365-113,779,787 with *p*-value = 7.81×10^−7^ overlapped the four eSCAN signals (**a**). Each of the four eSCAN signals is smaller in size than the SCANG detected regions (**b**). Specifically, eSCAN detected chr10:113,770,735-113,773,147 with *p*-value = 2.84×10^−6^; chr10:113,773,148-113,774,046 with *p*-value = 6.59×10^−6^; chr10:113,774,047-113,775,910 with *p*-value = 9.55×10^−6^ and chr10:113,775,911-113,778,291 with *p*-value = 1.92×10^−5^). Each SCANG signal contains two eSCAN signals (**c**). We then removed the associated variants in the overlapped regions between eSCAN and SCANG (which are regions detected by eSCAN since in both cases, the eSCAN regions are subsets of the SCANG regions), and re-did SCANG analysis using the retained variants only. Both regions then became insignificant (*p*-values > 0.02) using SCANG (**d**), suggesting that the sub-regions detected by eSCAN were most likely the functional regions contributing to the original significant signals.

The computational complexity of eSCAN depends on the sample size, the number of considered enhancers along a certain chromosome, and the number of rare variants residing in enhancer regions. For JHS (n=1,970) and WHI (n=10,727) eSCAN takes an average of 3h and 26h, respectively, to examine all the sets of rare variants along one chromosome, using our cluster computing platform with one computing node and 8Gb of memory (**Fig. S8**) while SCANG limited to enhancer regions takes an average of 2.6 days and 5.3 days respectively as more eigen-decomposition steps are performed.

To justify the choice of the number of PCs, we additionally applied eSCAN to HGB in the WHI cohort, using the first 5 and 8 PCs respectively. As shown in **Fig. S9**, the number of PCs doesn’t impact the results since the *p*-values are very well consistent. To justify the choice of SCANG window size range, we ran two more SCANG applications on the WHI cohort to analyze HGB using two different sets of window sizes: (1) L_min=90, L_max = 220; and (2) L_min=140, L_max = 250. The mirror Manhattan plots (**Fig. S10**) show similar pattern at all sets of window sizes, suggesting that the range of window sizes did not substantially affect SCANG’s results.

## Discussion

We propose here eSCAN, a novel aggregation method for whole genome sequencing analysis, which can integrate various types of functional information to aggregate enhancers or putative regulatory regions from WGS data and test for association with phenotypes of interest. Our method has several important advantages: (1) it has higher power and lower false positive rate, enabling it to accurately detect more significant signals than other methods (**Fig. 2; Fig. S2-5**); (2) the signals identified by eSCAN are of shorter sizes, which suggests eSCAN can more accurately locate the associated variants; (3) eSCAN boosts the biological interpretation of detected signals by incorporating functional annotation; (4) it is computationally efficient (**Fig. S8**).

The reason that eSCAN can improve the power for detecting causal variants is that eSCAN restricts the aggregation test only to variants from putative enhancer regions, where the causal variants are mostly likely to reside. In contrast, SCANG simply assigns variants into fixed-size sliding windows though the window length may change within a pre-specified range, which may identify variants from regions encompassing multiple enhancer regions with distinct functions as well as non-regulatory regions, thus leading to a lower power and a higher false positive rate in a fine-mapping sense. Hence, compared to SCANG, eSCAN is more likely to better pinpoint the regions with the causal variants. Furthermore, eSCAN performs a scanning procedure within each significant enhancer to further narrow down the potential causal region, attaining higher-resolution fine-mapping.

eSCAN can be viewed as an extension of SCANG with respect to its use of dynamic searching windows and use of the *p*-value as its test statistic (4). But it differs from SCANG in several key ways. SCANG restricts the size of searching windows within a pre-specified range and then tests all possible windows, “randomly” identifying some large regions across the genome regardless of their biological functions. eSCAN allows more flexible and biologically meaningful searching windows. It aggregates variants in putative enhancer regions to perform test within each enhancer (**Fig. 1**). In addition, eSCAN builds on fastSKAT, a computationally efficient approach to approximate the null distribution of SKAT statistics (11). We adopt an omnibus test that uses the aggregated Cauchy method via ACAT to combine *p*-values from fastSKAT using two different weights, Beta(1, 1) and Beta(1, 25). This omnibus test can additionally include burden test if desired. Compared to the optimal test in SCANG, the omnibus test has two advantages: 1) ACAT is flexible and can accommodate different choices of weights but the optimal test is not able to combine *p*-values under different weights; and 2) ACAT method is computationally more efficient and thus more scalable than the optimal test.

Based on our simulations in a variety of scenarios, eSCAN can be flexibly applied to different phenotypes, both quantitative and qualitative, and is able to detect more significant signals than competing methods with a better control over false positive rate than other WGS based methods (**Fig. 2; Fig. S2-4**). Using WGS data from the JHS and WHI studies, we demonstrate an enrichment of association signals using eSCAN procedure. It can detect reported signals which are not found by SCANG procedures, indicating that it is less likely to miss important regions. In addition, the regions detected by eSCAN are of shorter size than those of SCANG on average. By removing eSCAN signals from WGS data on chromosome 10 and re-running SCANG procedures, we verify that, at least for this segment, the signals detected by eSCAN drive the significant associations in larger regions identified by SCANG (**Fig. 3; Fig. S5**), a pattern we anticipate would be true for many associated regions.

Despite the modest sample size available for our blood cell trait analysis, interesting and biologically plausible rare and low frequency variant enhancer region signals were identified in our analyses from WHI. Of the genes regulated by replicated regions, *BACH2* (regulated by a region on chr6:90,423,754-90,425,200) is a key immune cell regulatory factor and is crucial for the maintenance of regulatory T-cell function and B-cell maturation (27). Among other interesting genes *CCL18* (regulated by a region on chr17:35,982,416-35,983,367, which was not replicated in JHS) was reported to stimulate the bone marrow overall, which could lead to increased platelets(28). These findings suggest that the associated enhancer regions identified by eSCAN may in fact play key regulatory relevant to the biological functions of blood cells, with eSCAN finding regions were not identified using the SCANG method. We do note, however, that these findings should be considered preliminary, given our modest sample size, and could be influenced by unadjusted for selection bias in WHI TOPMed sampling (enrichment for stroke and venous thromboembolism) and lack of adjustment for a genetic relationship matrix which could better capture cryptic relatedness and differential ancestry unadjusted for by PCs. However, these issues impact eSCAN and SCANG equally, and do not change our central methods comparison findings.

With respect to the weights in fastSKAT, we used two standard MAF-based weights: one is the Beta distribution with *a*_1_ = *a*_2_ =1 reflecting that all the variants have equal effect size, the other is *a*_1_=*a*_2_ =1, 25 upweighting rarer variants. One can also use external measures by incorporating individual level functional annotations, such as FATHMM-XF (29) and STAAR (30), as the weight for each variant. Incorporation of functional evidence has demonstrated its values in variant level association studies (31,32). In addition, the eSCAN framework is flexible regarding its unit aggregate tests. In our implementation, we use fastSKAT because of its small computational cost, but other aggregate tests can also be used, such as SMMAT, a recently proposed test which is an efficient variant set mixed model association test (33).

Another attractive feature of eSCAN is its significance threshold. Since candidate regions are highly likely to be correlated because of either physical overlapping or LD, making the set-based *p*-values also correlated, the classic Bonferroni correction would be too conservative. While we do use a classic Bonferroni correction in our real data example from WHI, due to the small sample size available to us for replication, this is almost certainly over-conservative. eSCAN provides two estimations of significance threshold, either empirically or analytically, using the strategies from SCANG and WGScan respectively, which have demonstrated significant enrichments of signals in Li *et al*. (4) and He *et al*. (10). In addition, although our analyses focused on unrelated individuals, it can be readily extended to related samples by replacing the generalized linear model (GLM) with the generalized linear mixed model (GLMM) in the first step (4).

One potential limitation of eSCAN is the lack of base pair resolution in defining regions important for gene regulation, due to the sparsity of reads with most Hi-C and chromatin conformation assays (leading to resolution as broad as 40 kb when assessing interactions between genomic regions). ATAC-seq data, albeit much finer resolution, still results in open chromatin peak regions that usually contain multiple rare variants, particularly as sample size increases, hurdling inference at the resolution of single base pair or single variant. These limitations are intrinsic to the functional annotation data employed rather than to the eSCAN methodology. We anticipate that rapid technological improvements in the functional annotation datasets will continue mitigating these issues by providing increasingly finer resolution and more comprehensive data, which would render eSCAN even more valuable in the near future.

## Data and Code Availability

This paper did not generate any datasets. TOPMed data from the Women’s Health Initiative is available to approved researchers through dbGaP (phs001237), with phenotype data available at phs000200. TOPMed data from the Jackson Heart Study Data is also available to approved researchers through dbGaP (phs000964), with phenotype data available at phs000286. Data is also available with an approved manuscript proposal through https://www.jacksonheartstudy.org/ (JHS) and https://www.whi.org/ (WHI).

## Web Resources

We developed an R package for the eSCAN procedure. The package is available at https://github.com/yingxi-kaylee/eSCAN.

## Acknowledgements

We thank Zilin Li for helpful input on SCANG methods and implementation.

The Jackson Heart Study (JHS) is supported and conducted in collaboration with Jackson State University (HHSN268201800013I), Tougaloo College (HHSN268201800014I), the Mississippi State Department of Health (HHSN268201800015I/HHSN26800001) and the University of Mississippi Medical Center (HHSN268201800010I, HHSN268201800011I and HHSN268201800012I) contracts from the National Heart, Lung, and Blood Institute (NHLBI) and the National Institute for Minority Health and Health Disparities (NIMHD). The authors also wish to thank the staff and participants of the JHS.

The WHI program is funded by the National Heart, Lung, and Blood Institute, National Institutes of Health, U.S. Department of Health and Human Services through contracts HHSN268201600018C, HHSN268201600001C, HHSN268201600002C, HHSN268201600003C, and HHSN268201600004C. The authors thank the WHI investigators and staff for their dedication, and the study participants for making the program possible. A listing of WHI investigators can be found at: https://www-whi-org.s3.us-west-2.amazonaws.com/wp-content/uploads/WHI-Investigator-Short-List.pdf.

The views expressed in this manuscript are those of the authors and do not necessarily represent the views of the National Heart, Lung, and Blood Institute; the National Institutes of Health; or the U.S. Department of Health and Human Services.

Molecular data for the Trans-Omics in Precision Medicine (TOPMed) program was supported by the National Heart, Lung and Blood Institute (NHLBI). Genome sequencing for “NHLBI TOPMed: The Jackson Heart Study” (phs000964.v1.p1) was performed at the Northwest Genomics Center (HHSN268201100037C). Genome sequencing for “NHLBI TOPMed: Women’s Health Initiative (WHI)” (phs001237) was performed by Broad Genomics (HHSN268201500014C). Core support including centralized genomic read mapping and genotype calling, along with variant quality metrics and filtering were provided by the TOPMed Informatics Research Center (3R01HL-117626-02S1; contract HHSN268201800002I). Core support including phenotype harmonization, data management, sample-identity QC, and general program coordination were provided by the TOPMed Data Coordinating Center (R01HL-120393; U01HL-120393; contract HHSN268201800001I). We gratefully acknowledge the studies and participants who provided biological samples and data for TOPMed. A list of TOPMed investigators represented by the TOPMed banner can be found at https://www.nhlbiwgs.org/topmed-banner-authorship.

## REFERENCES

1. Morrison, A.C., Huang, Z., Yu, B., Metcalf, G., Liu, X., Ballantyne, C., Coresh, J., Yu, F., Muzny, D., Feofanova, E. et al. (2017) Practical Approaches for Whole-Genome Sequence Analysis of Heart- and Blood-Related Traits. Am J Hum Genet, 100, 205–215.

2. Morrison, A.C., Voorman, A., Johnson, A.D., Liu, X., Yu, J., Li, A., Muzny, D., Yu, F., Rice, K., Zhu, C. et al. (2013) Whole-genome sequence-based analysis of high-density lipoprotein cholesterol. Nat Genet, 45, 899–901.

3. Natarajan, P., Peloso, G.M., Zekavat, S.M., Montasser, M., Ganna, A., Chaffin, M., Khera, A.V., Zhou, W., Bloom, J.M., Engreitz, J.M. et al. (2018) Deep-coverage whole genome sequences and blood lipids among 16,324 individuals. Nature communications, 9, 3391.

4. Li, Z., Li, X., Liu, Y., Shen, J., Chen, H., Zhou, H., Morrison, A.C., Boerwinkle, E. and Lin, X. (2019) Dynamic Scan Procedure for Detecting Rare-Variant Association Regions in Whole-Genome Sequencing Studies. Am J Hum Genet, 104, 802–814.

5. Wu, M., Lee, S., Cai, T., Li, Y., Boehnke, M. and Lin, X. (2011) Rare-variant association testing for sequencing data with the sequence kernel association test. American journal of human genetics, 89, 82–93.

6. Farh, K.K., Marson, A., Zhu, J., Kleinewietfeld, M., Housley, W.J., Beik, S., Shoresh, N., Whitton, H., Ryan, R.J., Shishkin, A.A. et al. (2015) Genetic and epigenetic fine mapping of causal autoimmune disease variants. Nature, 518, 337–343.

7. Onengut-Gumuscu, S., Chen, W.-M., Burren, O., Cooper, N.J., Quinlan, A.R., Mychaleckyj, J.C., Farber, E., Bonnie, J.K., Szpak, M., Schofield, E. et al. (2015) Fine mapping of type 1 diabetes susceptibility loci and evidence for colocalization of causal variants with lymphoid gene enhancers. Nature Genetics, 47, 381–386.

8. Gallagher, M.D. and Chen-Plotkin, A.S. (2018) The Post-GWAS Era: From Association to Function. Am J Hum Genet, 102, 717–730.

9. Wu, C. and Pan, W. (2018) Integration of Enhancer-Promoter Interactions with GWAS Summary Results Identifies Novel Schizophrenia-Associated Genes and Pathways. Genetics, 209, 699.

10. He, Z., Xu, B., Buxbaum, J. and Ionita-Laza, I. (2019) A genome-wide scan statistic framework for whole-genome sequence data analysis. Nature communications, 10, 3018.

11. Lumley, T., Brody, J., Peloso, G., Morrison, A. and Rice, K. (2018) FastSKAT: Sequence kernel association tests for very large sets of markers. Genet Epidemiol, 42, 516–527.

12. Astle, W.J., Elding, H., Jiang, T., Allen, D., Ruklisa, D., Mann, A.L., Mead, D., Bouman, H., Riveros-Mckay, F., Kostadima, M.A. et al. (2016) The Allelic Landscape of Human Blood Cell Trait Variation and Links to Common Complex Disease. Cell, 167, 1415-1429.e1419.

13. Liu, Y., Chen, S., Li, Z., Morrison, A.C., Boerwinkle, E. and Lin, X. (2019) ACAT: A Fast and Powerful p Value Combination Method for Rare-Variant Analysis in Sequencing Studies. Am J Hum Genet, 104, 410–421.

14. Taylor, H.A., Jr., Wilson, J.G., Jones, D.W., Sarpong, D.F., Srinivasan, A., Garrison, R.J., Nelson, C. and Wyatt, S.B. (2005) Toward resolution of cardiovascular health disparities in African Americans: design and methods of the Jackson Heart Study. Ethnicity & disease, 15, S6-4-17.

15. The Women’s Health Initiative Study Group. (1998) Design of the Women’s Health Initiative clinical trial and observational study. Controlled clinical trials, 19, 61–109.

16. Nalls, M.A., Wilson, J.G., Patterson, N.J., Tandon, A., Zmuda, J.M., Huntsman, S., Garcia, M., Hu, D., Li, R., Beamer, B.A. et al. (2008) Admixture mapping of white cell count: genetic locus responsible for lower white blood cell count in the Health ABC and Jackson Heart studies. Am J Hum Genet, 82, 81–87.

17. Reich, D., Nalls, M.A., Kao, W.H., Akylbekova, E.L., Tandon, A., Patterson, N., Mullikin, J., Hsueh, W.C., Cheng, C.Y., Coresh, J. et al. (2009) Reduced neutrophil count in people of African descent is due to a regulatory variant in the Duffy antigen receptor for chemokines gene. PLoS Genet, 5, e1000360.

18. Chen, M.H., Raffield, L.M., Mousas, A., Sakaue, S., Huffman, J.E., Moscati, A., Trivedi, B., Jiang, T., Akbari, P., Vuckovic, D. et al. (2020) Trans-ethnic and Ancestry-Specific Blood-Cell Genetics in 746,667 Individuals from 5 Global Populations. Cell, 182, 1198-1213.e1114.

19. Javierre, B.M., Burren, O.S., Wilder, S.P., Kreuzhuber, R., Hill, S.M., Sewitz, S., Cairns, J., Wingett, S.W., Várnai, C., Thiecke, M.J. et al. (2016) Lineage-Specific Genome Architecture Links Enhancers and Non-coding Disease Variants to Target Gene Promoters. Cell, 167, 1369-1384.e1319.

20. Gieger, C., Radhakrishnan, A., Cvejic, A., Tang, W., Porcu, E., Pistis, G., Serbanovic-Canic, J., Elling, U., Goodall, A.H., Labrune, Y. et al. (2011) New gene functions in megakaryopoiesis and platelet formation. Nature, 480, 201–208.

21. Kanai, M., Akiyama, M., Takahashi, A., Matoba, N., Momozawa, Y., Ikeda, M., Iwata, N., Ikegawa, S., Hirata, M., Matsuda, K. et al. (2018) Genetic analysis of quantitative traits in the Japanese population links cell types to complex human diseases. Nat Genet, 50, 390–400.

22. Kanai, M., Akiyama, M., Takahashi, A., Matoba, N., Momozawa, Y., Ikeda, M., Iwata, N., Ikegawa, S., Hirata, M., Matsuda, K. et al. (2018) Genetic analysis of quantitative traits in the Japanese population links cell types to complex human diseases. Nat Genet, 50, 390–400.

23. Pankratz, S., Bittner, S., Kehrel, B.E., Langer, H.F., Kleinschnitz, C., Meuth, S.G. and Gobel, K. (2016) The Inflammatory Role of Platelets: Translational Insights from Experimental Studies of Autoimmune Disorders. Int J Mol Sci, 17.

24. van der Harst, P., Zhang, W., Mateo Leach, I., Rendon, A., Verweij, N., Sehmi, J., Paul, D.S., Elling, U., Allayee, H., Li, X. et al. (2012) Seventy-five genetic loci influencing the human red blood cell. Nature, 492, 369–375.

25. Vuckovic, D., Bao, E.L., Akbari, P., Lareau, C.A., Mousas, A., Jiang, T., Chen, M.H., Raffield, L.M., Tardaguila, M., Huffman, J.E. et al. (2020) The Polygenic and Monogenic Basis of Blood Traits and Diseases. Cell, 182, 1214-1231.e1211.

26. Mousas, A., Ntritsos, G., Chen, M.-H., Song, C., Huffman, J., Tzoulaki, I., Elliott, P., Psaty, B., Auer, P., Johnson, A. et al. (2017) Rare coding variants pinpoint genes that control human hematological traits. PLoS genetics, 13, e1006925.

27. Afzali, B., Grönholm, J., Vandrovcova, J., O’Brien, C., Sun, H.W., Vanderleyden, I., Davis, F.P., Khoder, A., Zhang, Y., Hegazy, A.N. et al. (2017) BACH2 immunodeficiency illustrates an association between super-enhancers and haploinsufficiency. Nature immunology, 18, 813–823.

28. Wimmer, A., Khaldoyanidi, S.K., Judex, M., Serobyan, N., DiScipio, R.G. and Schraufstatter, I.U. (2006) CCL18/PARC stimulates hematopoiesis in long-term bone marrow cultures indirectly through its effect on monocytes. Blood, 108, 3722–3729.

29. Rogers, M.F., Shihab, H.A., Mort, M., Cooper, D.N., Gaunt, T.R. and Campbell, C. (2018) FATHMM-XF: accurate prediction of pathogenic point mutations via extended features. Bioinformatics, 34, 511–513.

30. Li, X., Li, Z., Zhou, H., Gaynor, S.M., Liu, Y., Chen, H., Sun, R., Dey, R., Arnett, D.K., Aslibekyan, S. et al. (2020) Dynamic incorporation of multiple in silico functional annotations empowers rare variant association analysis of large whole-genome sequencing studies at scale. Nat Genet, 52, 969–983.

31. Pickrell, J.K. (2014) Joint analysis of functional genomic data and genome-wide association studies of 18 human traits. Am J Hum Genet, 94, 559–573.

32. Yang, J., Fritsche, L.G., Zhou, X. and Abecasis, G. (2017) A Scalable Bayesian Method for Integrating Functional Information in Genome-wide Association Studies. Am J Hum Genet, 101, 404–416.

33. Chen, H., Huffman, J.E., Brody, J.A., Wang, C., Lee, S., Li, Z., Gogarten, S.M., Sofer, T., Bielak, L.F., Bis, J.C. et al. (2019) Efficient Variant Set Mixed Model Association Tests for Continuous and Binary Traits in Large-Scale Whole-Genome Sequencing Studies. Am J Hum Genet, 104, 260–274.

