## Supplemental Figures and Table for "eSCAN: Scan Regulatory Regions for Aggregate Association Testing using Whole Genome Sequencing Data"

### **Supplemental Materials**

#### Supplemental Figures

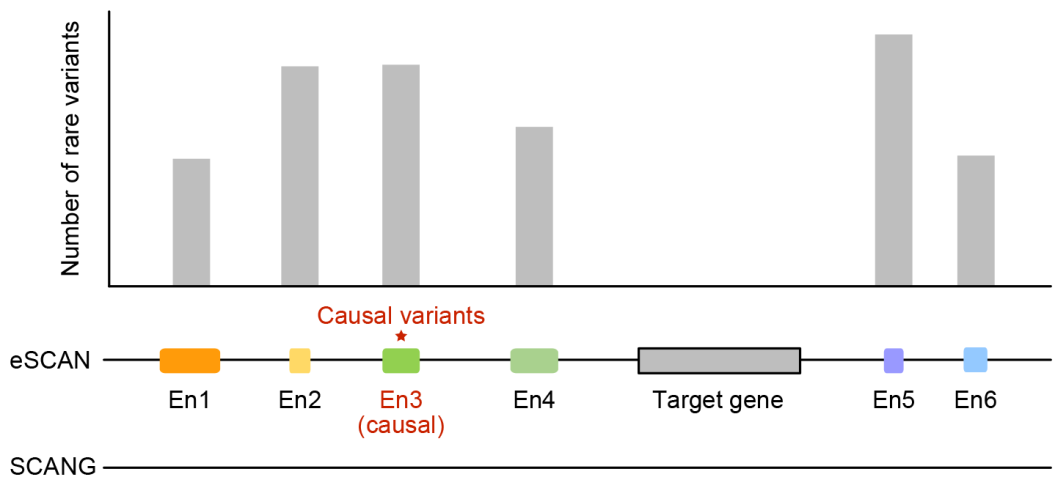

**Figure S1. An illustration of simulation framework.**

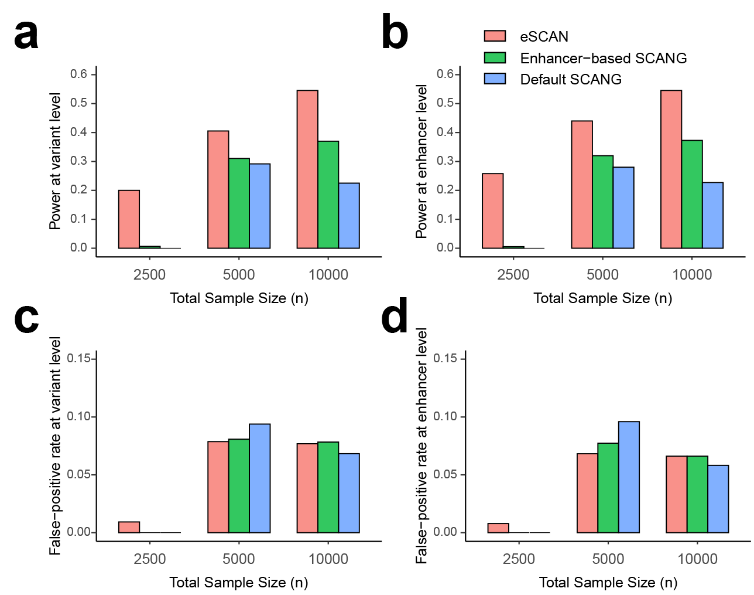

**Figure S2. Power and false positive rate comparison of eSCAN and SCANG for dichotomous outcome at various sample sizes.** We evaluated the performance of eSCAN for dichotomous outcome at various sample sizes. The total sample sizes considered were 2,500, 5,000 and 10,000. For each configuration, we compared three methods: eSCAN and two versions of SCANGs: enhancer-based SCANG (aggregating enhancers across the genome) and default SCANG (scan the whole genome). We evaluated power at both variant and enhancer level, by causal variant detection rate and causal enhancer detection rate, respectively. Also, we evaluated false positive rate at both variant and enhancer level. All criteria were calculated at the genome-wide type I error rate $\alpha=0.05$. We simulated 10% of enhancers across the genome to be causal; within each causal enhancer, we randomly selected 20% of variants with MAF$<5\%$ to be causal ones; the effect size of the causal variants was a decreasing function of MAF, $\beta=c|log_{10}MAF|$, where $c=0.255$. **a.** Power at variant level (a.k.a. causal variant detection rate). **b.** Power at enhancer level (a.k.a. causal enhancer detection rate). **c.** False positive rate at variant level. **d.** False positive rate at enhancer level.

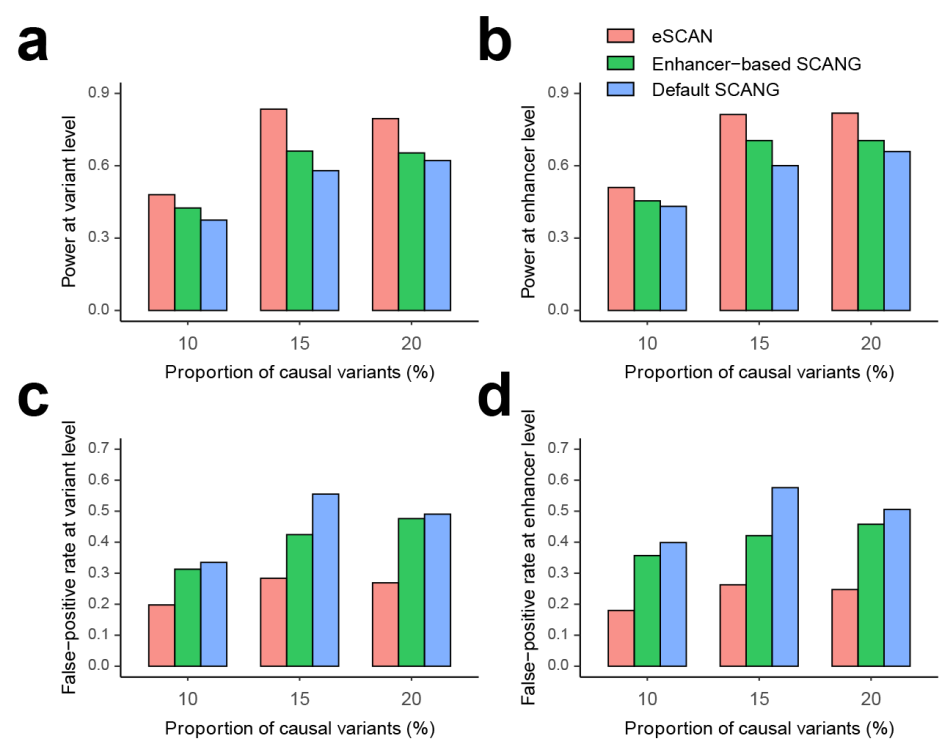

**Figure S3. Power and false positive rate comparison of eSCAN and SCANG for continuous outcome at various portions of causal enhancers.** We evaluated the performance of eSCAN for continuous outcome at various portions of causal enhancers across the whole genome. The portions considered were 5%, 10% and 15%. For each configuration, we compared three methods: eSCAN and two versions of SCANGs: enhancer-based SCANG (aggregating enhancers across the genome) and default SCANG (scan the whole genome). We evaluated power at variant and enhancer level, by causal variant detection rate and causal enhancer detection rate, respectively. Also, we evaluated false positive rate at variant and enhancer level. All the criteria were calculated at the genome-wide type I error rate $\alpha=0.05$. We randomly selected 20% of variants with MAF$<5\%$ to be causal ones, within each causal enhancer in the simulated 10,000 whole-genome data; the effect size of the causal variants was a decreasing function of MAF, $\beta=c|log_{10}MAF|$, where $c=0.18$. **a.** Power at variant level (a.k.a. causal variant detection rate). **b.** Power at enhancer level (a.k.a. causal-enhancer detection rate). **c.** False positive rate at variant level. **d.** False positive rate at enhancer level.

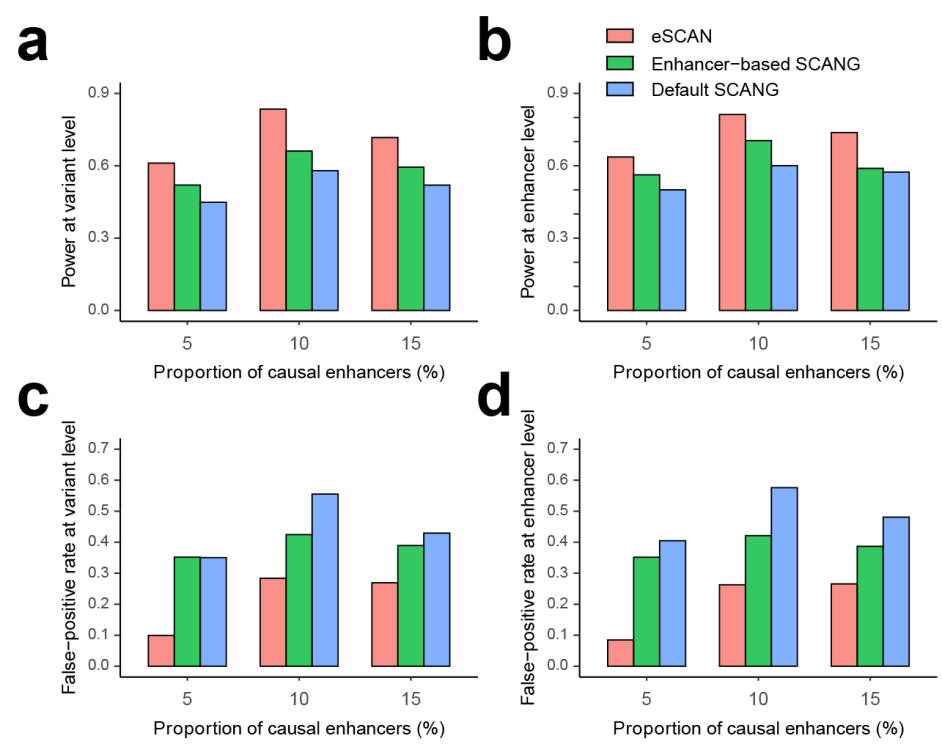

**Figure S4. Power and false positive rate comparison of eSCAN and SCANG for continuous outcome at various portions of causal variants.** We evaluated the performance of eSCAN for continuous outcome as the various portions of causal rare variants. The considered portions of causal variants with MAF$<5\%$ within causal enhancers were 10%, 15% and 20%. For each configuration, we compared three methods: eSCAN and two versions of SCANGs: enhancer-based SCANG (aggregating enhancers across the genome) and default SCANG (scan the whole genome). We evaluated power at variant and enhancer level, by causal variant detection rate and causal enhancer detection rate, respectively. Also, we evaluated false positive rate at variant and enhancer level. All the criteria were calculated at the genome-wide type I error rate $\alpha=0.05$. For each setting, in the simulated 10,000 whole-genome data, we simulated 10% of enhancers across the genome to be causal, then randomly selected the aforementioned portion of variants to be causal ones; the effect size of the causal variants was a decreasing function of MAF, $\beta=c|log_{10}MAF|$, where $c=0.18$. **a.** Power at variant level (a.k.a. causal variant detection rate). **b.** Power at enhancer level (a.k.a. causal enhancer detection rate). **c.** False positive rate at variant level. **d.** False positive rate at enhancer level.

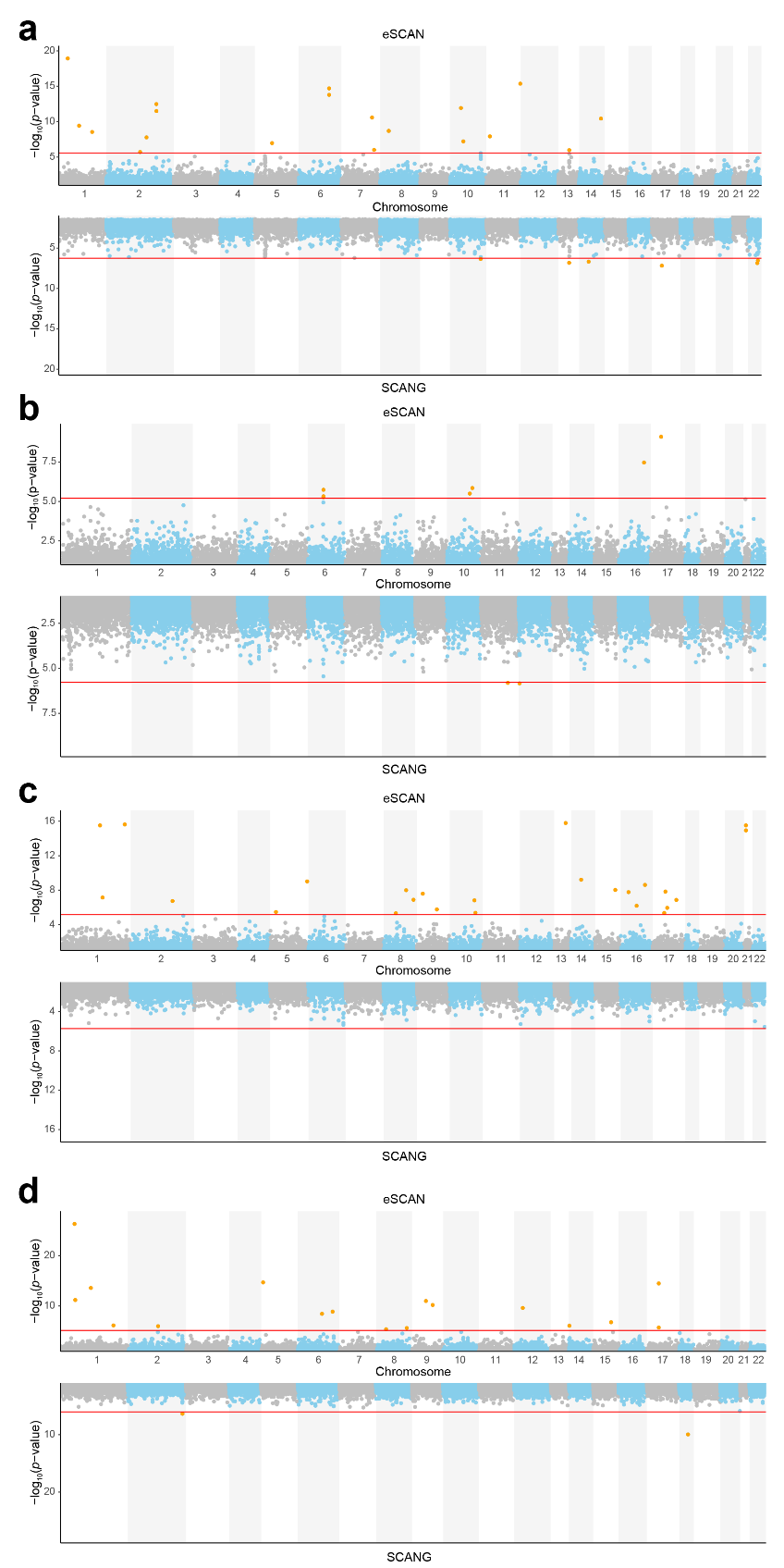

**Figure S5. Mirror Manhattan plots of associations of genome-wide regions with four blood cell traits in the WHI study.** **a.** A mirror Manhattan plot for white blood cell count (WBC). **b.** A mirror Manhattan plot for hematocrit (HCT). **c.** A mirror Manhattan plot for hemoglobin (HGB). **d.** A mirror Manhattan plot for platelet count (PLT).

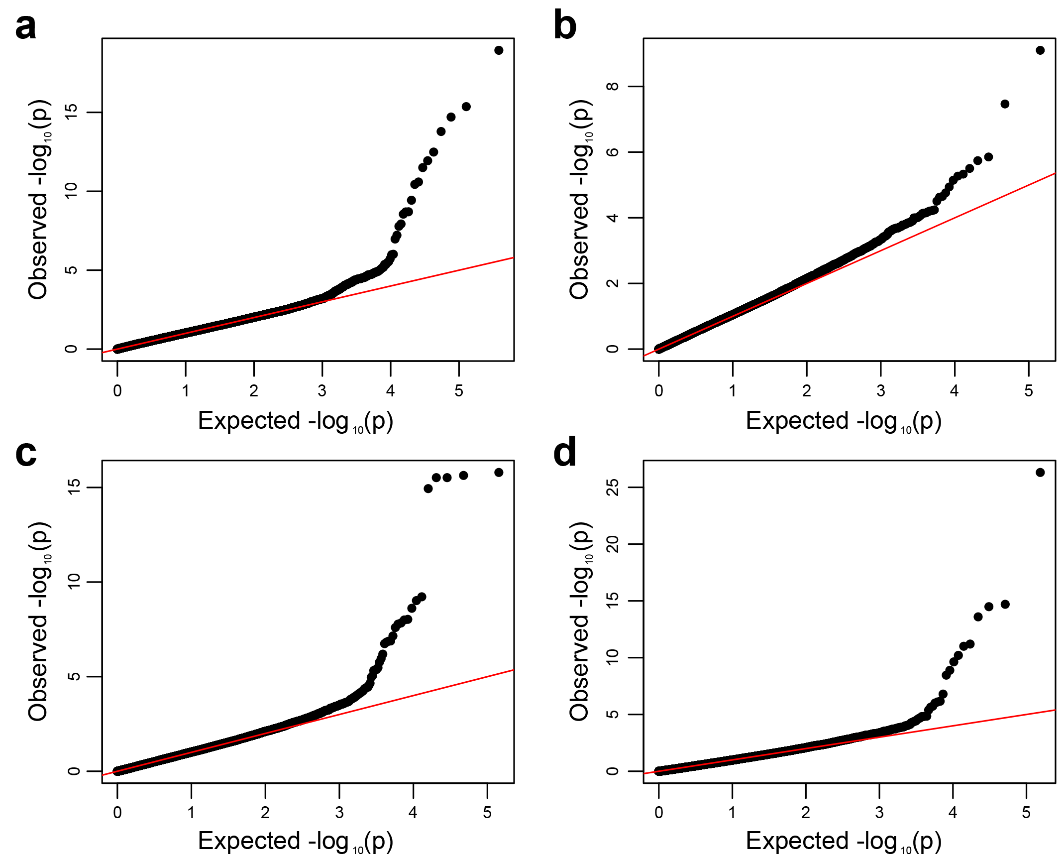

**Figure S6. QQ plots of associations of genome-wide regions with four blood cell traits in the WHI study.** QQ plot for white blood cell count (WBC) with genomic control lambda = 1.14 (**a**), hematocrit (HCT) with lambda = 1.00 (**b**), hemoglobin (HGB) with lambda = 1.00 (**c**), and platelet count (PLT) with lambda = 0.93 (**d**).

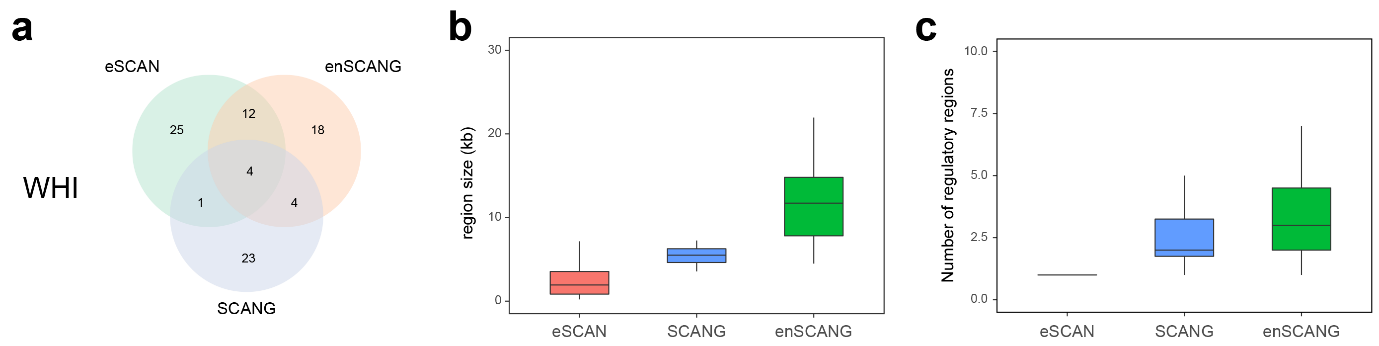

**Figure S7. Results using empirical threshold by analysing TOPMed WBC data. a.** A Venn diagram of significant regions identified by three methods, eSCAN, enhancer-based SCANG and the default SCANG, in the WHI discovery samples. Enhancer-based SCANG (“enSCANG” in the figure) refers to applying SCANG to the subset of rare variants falling into any enhancer region as defined using PC-HiC annotation, whereas “SCANG” in the figure indicates the default procedure of testing all rare variants. **b.** The length of identified regions by the three methods. **c.** The number of regulatory elements included in the identified regions by the three methods.

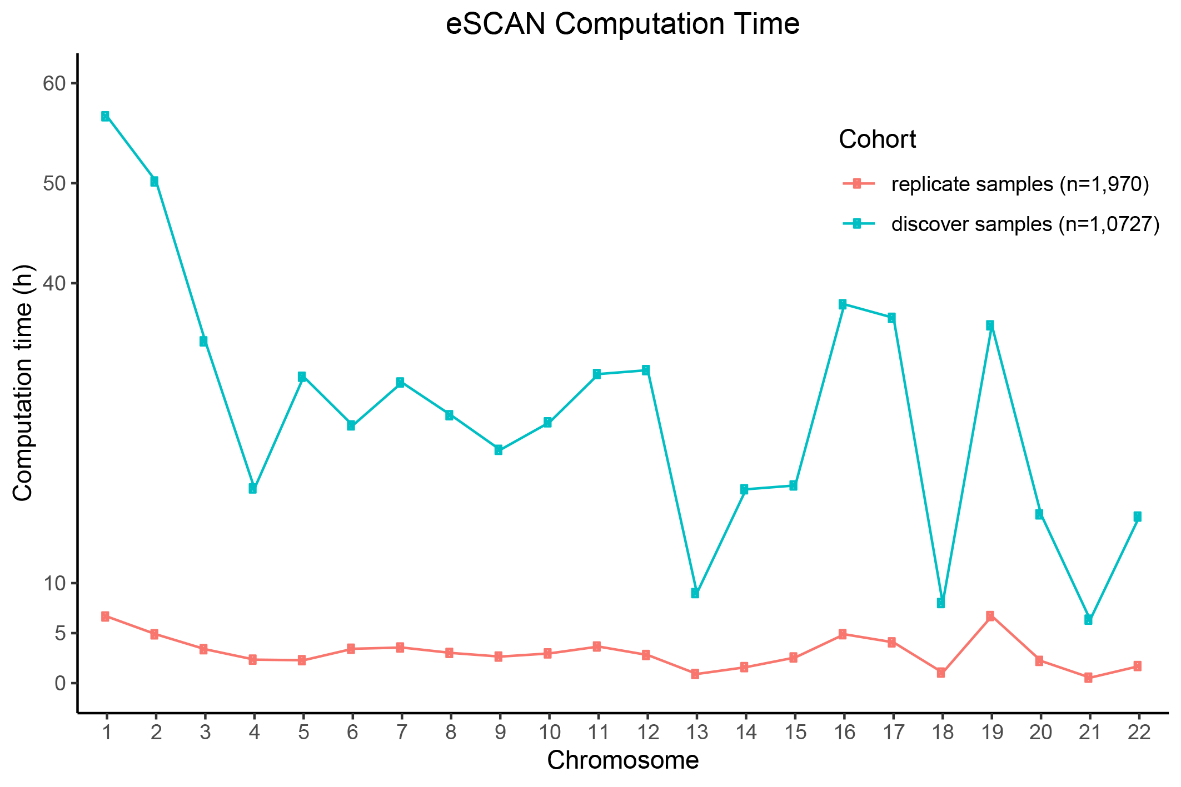

**Figure S8. Computational time of eSCAN.**

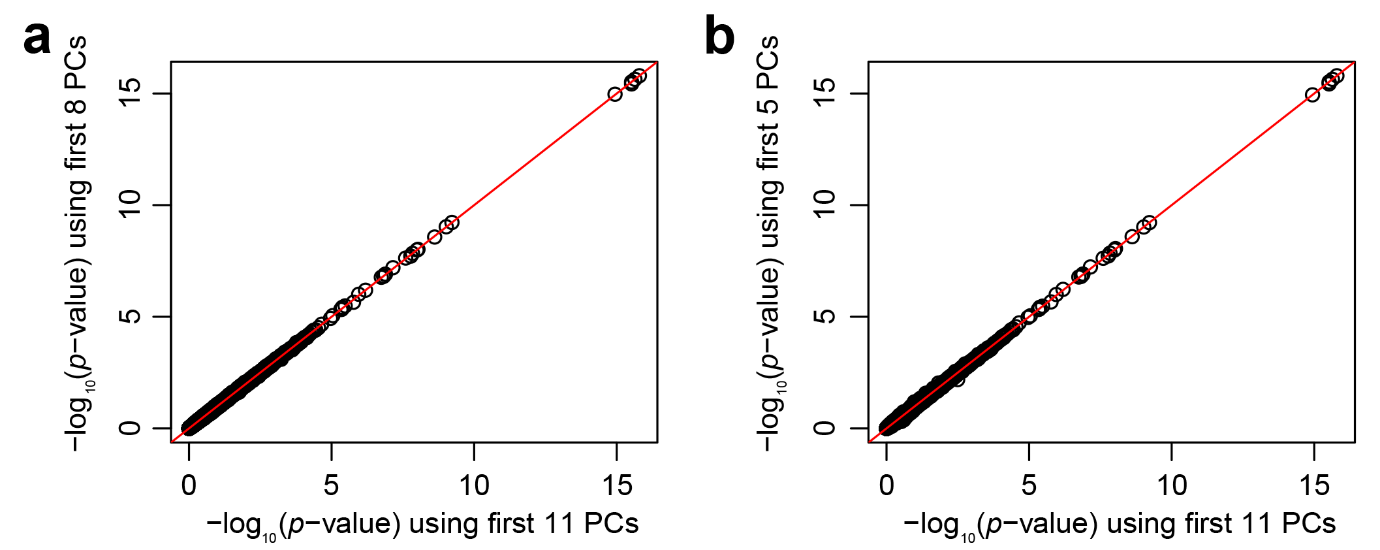

**Figure S9. Scatter plots of –log_10_(*p*-value) using different number of principle components (PCs) for hemoglobin (HGB) in the WHI study. a.** Scatter plot of –log_10_(*p*-value) using the first 11 PCs (x-axis) and 8 PCs (y-axis) respectively to analyse HGB in the WHI study. **b.** Scatter plot of –log_10_(*p*-value) using the first 11 PCs (x-axis) and 5 PCs (y-axis) respectively to analyse HGB in the WHI study.

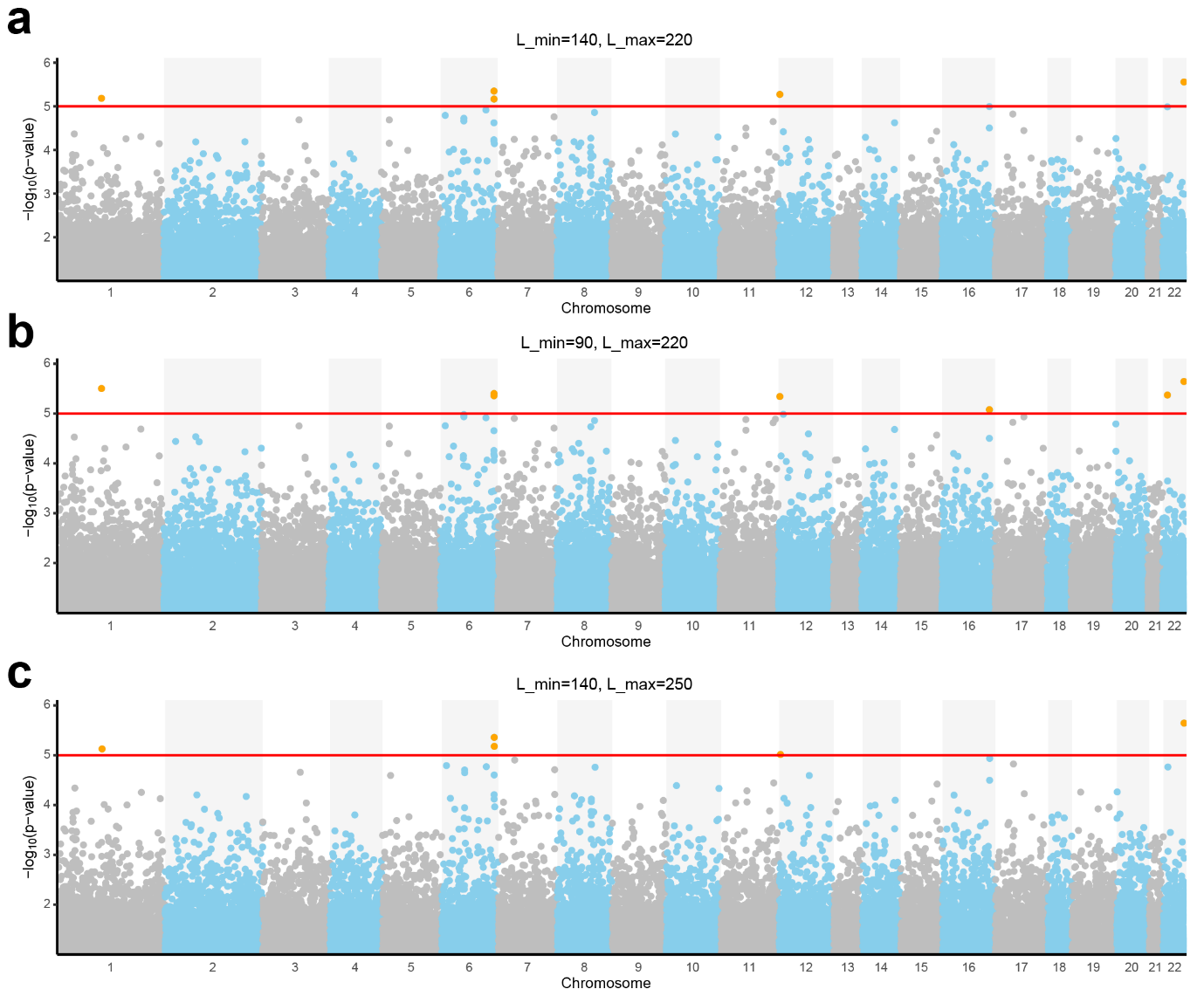

**Figure S10. Mirror Manhattan plots using different parameters (‘Lmin’ and ‘Lmax’) in SCANG for hemoglobin (HGB) in the WHI study.** We additionally ran two more sets of SCANG in the WHI cohort to analyze HGB. We change the two parameters Lmin and Lmax one at a time: (1) we keep the Lmax to 220 and lower the Lmin to 90; (2) we keep the Lmin to 140 and increase the Lmax to 250. **a.** A mirror Manhattan plot for Lmin=140, Lmax=220 and Lmin=90, Lmax=220. **b.** A mirror Manhattan plot for Lmin=140, Lmax=220 and Lmin=140, Lmax=250.

#### Supplemental Tables

**Table S1. Study-specific descriptive statistics of participating cohorts (mean (SD)).** HGB, hemoglobin; HCT, hematocrit; PLT, platelet count; WBC, white blood cell count.

|  | **Age** | **%Female** | **WBC (10^9/L)** | **HCT (%)** | **HGB (g/dL)** | **PLT (10^9/L)** | **Sample Size** |
| --- | --- | --- | --- | --- | --- | --- | --- |
| Discovery (WHI) | 66.78 (6.78) | 100 | 6.12 (2.21) | 40.21 (2.96) | 13.52 (1.04) | 249.98 (63.82) | 10727 |
| Replication (JHS) | 55.01 (12.26) | 60.94 | 5.60 (1.80) | 39.45 (4.22) | 13.09 (1.47) | 246.80 (61.17) | 1970 |

**Table S2. Significant results using Bonferroni correction for enhancer-based SCANG for blood cell traits in TOPMed WGS data.** Chr, chromosome; Start, start position (hg38); End, end position (hg38); Trait (PLT, platelet count; WBC, white blood cell count).

| Chr | Start | End | *P*-value | Trait |
| --- | --- | --- | --- | --- |
| 18 | 54,746,727 | 54,776,771 | 1.09E-10 | PLT |
| 10 | 113,755,577 | 113,783,415 | 4.38E-07 | WBC |
| 13 | 52,011,368 | 52,020,214 | 1.49E-07 | WBC |
| 14 | 63,429,139 | 63,438,028 | 2.05E-07 | WBC |
| 17 | 37,709,185 | 37,716,290 | 6.56E-08 | WBC |
| 22 | 40,921,264 | 40,943,211 | 1.37E-07 | WBC |
| 22 | 41,292,371 | 41,304,107 | 2.82E-07 | WBC |

**Table S3. Significant results using Bonferroni correction for the default SCANG for white blood cell count (WBC) in TOPMed WGS data.** Chr, chromosome; Start, start position (hg38); End, end position (hg38).

| Chr | Start | End | *P*-value |
| --- | --- | --- | --- |
| 2 | 40,315,899 | 40,321,177 | 1.64E-07 |
| 4 | 7,230,720 | 7,234,650 | 6.25E-07 |
| 4 | 49,708,861 | 49,710,138 | 6.15E-07 |
| 4 | 49,710,772 | 49,714,339 | 9.85E-10 |
| 4 | 53,787,731 | 53,793,156 | 3.16E-07 |
| 4 | 66,919,127 | 66,926,378 | 2.97E-07 |
| 4 | 78,403,805 | 78,409,475 | 4.13E-07 |
| 5 | 65,940,607 | 65,944,162 | 5.67E-07 |
| 5 | 65,952,476 | 65,957,214 | 8.70E-07 |
| 5 | 66,022,589 | 66,028,304 | 8.83E-07 |
| 5 | 66,053,510 | 66,058,768 | 4.92E-07 |
| 5 | 66,125,067 | 66,131,052 | 3.50E-07 |
| 7 | 99,059,416 | 99,064,059 | 4.66E-07 |
| 8 | 1,732,213 | 1,735,219 | 8.60E-07 |
| 8 | 59,102,832 | 59,107,933 | 6.99E-07 |
| 8 | 97,605,029 | 97,611,256 | 6.52E-07 |
| 8 | 106,396,366 | 106,403,182 | 2.13E-07 |
| 8 | 129,688,805 | 129,695,356 | 5.37E-07 |
| 9 | 134,607,546 | 134,613,472 | 8.87E-07 |
| 10 | 98,051,840 | 98,057,262 | 3.04E-07 |
| 10 | 113,767,467 | 113,773,998 | 2.66E-07 |
| 10 | 113,774,365 | 113,779,787 | 7.81E-07 |
| 12 | 94,158,340 | 94,164,463 | 1.68E-07 |
| 13 | 51,986,990 | 51,992,556 | 1.19E-07 |
| 13 | 52,011,726 | 52,018,111 | 2.34E-07 |
| 13 | 52,035,240 | 52,042,482 | 2.42E-08 |
| 13 | 52,087,348 | 52,094,303 | 2.37E-07 |
| 13 | 52,095,491 | 52,101,113 | 3.83E-08 |
| 13 | 52,113,234 | 52,120,333 | 8.74E-10 |
| 13 | 53,112,008 | 53,117,370 | 5.64E-10 |
| 13 | 53,128,179 | 53,134,339 | 3.44E-07 |
| 14 | 48,146,174 | 48,150,737 | 1.68E-07 |
| 14 | 105,863,207 | 105,864,144 | 3.89E-10 |
| 17 | 13,020,239 | 13,024,546 | 3.22E-07 |
| 17 | 37,710,209 | 37,716,290 | 2.25E-07 |
| 22 | 41,307,767 | 41,312,193 | 3.21E-07 |

**Table S4. Known GWAS loci within +/- 500kb of eSCAN detected regions.** Chr, chromosome; Start, start position (hg38) for eSCAN identified region; End, end position (hg38) for eSCAN identified region; *P*-value, *p*-value in the discovery samples; *P*-value (rep.), *p*-value in the replication samples; Trait (HGB, hemoglobin; PLT, platelet count; WBC, white blood cell count); SNP (rsID (reference) of nearby GWAS locus); Pos., position (hg38).

| **Chr** | **Start** | **End** | ***P*-value** | ***P*-value (rep.)** | **Cond. P-value** | **Trait** | **SNP** | **Pos.** |
| --- | --- | --- | --- | --- | --- | --- | --- | --- |
| 1 | 35,978,873 | 35,979,878 | 1.17E-19 | 0.11 | 3.63E-17 | WBC | rs3917932 (1-3) | 36,478,315 |
| 1 | 35,978,873 | 35,979,878 | 1.17E-19 | 0.11 | 3.63E-17 | WBC | rs148916169 (4) | 36,466,862 |
| 1 | 35,978,873 | 35,979,878 | 1.17E-19 | 0.11 | 3.63E-17 | WBC | rs3917932 (5) | 36,478,315 |
| 1 | 166,941,439 | 166,942,948 | 2.81E-09 | 0.34 | 1.03E-10 | WBC | rs864537 (6) | 167,442,147 |
| 9 | 32,397,109 | 32,404,847 | 2.52E-08 | 0.42 | 2.42E-08 | HGB | rs7045087 (5) | 32,455,264 |
| 17 | 35,982,416 | 35,983,367 | 3.24E-15 | 0.12 | 3.12E-14 | PLT | rs7221322 (1) | 35,546,753 |
| 17 | 35,982,416 | 35,983,367 | 3.24E-15 | 0.12 | 3.12E-14 | PLT | rs8073060 (5) | 35,548,243 |
| 17 | 35,982,416 | 35,983,367 | 3.24E-15 | 0.12 | 3.12E-14 | PLT | rs10512472 (7) | 35,557,785 |
| 17 | 35,982,416 | 35,983,367 | 3.24E-15 | 0.12 | 3.12E-14 | PLT | rs9908158 (5) | 35,563,064 |
| 17 | 35,982,416 | 35,983,367 | 3.24E-15 | 0.12 | 3.12E-14 | PLT | rs201192867 (5) | 35,572,827 |
| 17 | 35,982,416 | 35,983,367 | 3.24E-15 | 0.12 | 3.12E-14 | PLT | rs11653357 (5) | 35,596,588 |

#### Supplemental References

1.	Kanai, M., Akiyama, M., Takahashi, A., Matoba, N., Momozawa, Y., Ikeda, M., Iwata, N., Ikegawa, S., Hirata, M., Matsuda, K. *et al.* (2018) Genetic analysis of quantitative traits in the Japanese population links cell types to complex human diseases. *Nat Genet*, **50**, 390-400.

2.	Chen, M.H., Raffield, L.M., Mousas, A., Sakaue, S., Huffman, J.E., Moscati, A., Trivedi, B., Jiang, T., Akbari, P., Vuckovic, D. *et al.* (2020) Trans-ethnic and Ancestry-Specific Blood-Cell Genetics in 746,667 Individuals from 5 Global Populations. *Cell*, **182**, 1198-1213.e1114.

3.	Vuckovic, D., Bao, E.L., Akbari, P., Lareau, C.A., Mousas, A., Jiang, T., Chen, M.H., Raffield, L.M., Tardaguila, M., Huffman, J.E. *et al.* (2020) The Polygenic and Monogenic Basis of Blood Traits and Diseases. *Cell*, **182**, 1214-1231.e1211.

4.	Mousas, A., Ntritsos, G., Chen, M.-H., Song, C., Huffman, J., Tzoulaki, I., Elliott, P., Psaty, B., Auer, P., Johnson, A. *et al.* (2017) Rare coding variants pinpoint genes that control human hematological traits. *PLoS genetics*, **13**, e1006925.

5.	Astle, W.J., Elding, H., Jiang, T., Allen, D., Ruklisa, D., Mann, A.L., Mead, D., Bouman, H., Riveros-Mckay, F., Kostadima, M.A. *et al.* (2016) The Allelic Landscape of Human Blood Cell Trait Variation and Links to Common Complex Disease. *Cell*, **167**, 1415-1429.e1419.

6.	Pankratz, S., Bittner, S., Kehrel, B.E., Langer, H.F., Kleinschnitz, C., Meuth, S.G. and Gobel, K. (2016) The Inflammatory Role of Platelets: Translational Insights from Experimental Studies of Autoimmune Disorders. *Int J Mol Sci*, **17**.

7.	Gieger, C., Radhakrishnan, A., Cvejic, A., Tang, W., Porcu, E., Pistis, G., Serbanovic-Canic, J., Elling, U., Goodall, A.H., Labrune, Y. *et al.* (2011) New gene functions in megakaryopoiesis and platelet formation. *Nature*, 10.1038/nature10659.
